# Evaluation of object detection neural networks to find topologically associated domains in bulk Hi-C data

**DOI:** 10.64898/2026.10.20.760189

**Authors:** Niklas Philipp, Christopher Leon Grünnagel, Stefan Janssen, Cornelia Sigges

## Abstract

Topologically associated domains (TADs) shape the three-dimensional structure of chromatin and contribute to gene regulation by arranging regulatory elements and genes in spatial proximity. This spatial proximity is captured by Hi-C, an experimental method, which quantifies contact frequencies between all pairs of genomic loci. In bulk Hi-C data, TADs can be found in the form of gapped, nested and partially overlapping squares centered on the main diagonal of the Hi-C matrix. We shift the problem of TAD identification into the domain of computer vision by detecting TADs in Hi-C in the form of squares in images. For this we use object detection, which aims to classify object instances in images. This study evaluates the applicability of the object detection neural networks YOLOv8 and Mask R-CNN on the identification of TADs in bulk Hi-C data. We build image-like data from Hi-C matrices, predict reference TADs with a state-of-the-art TAD caller, train the models and evaluate the detections. Our results confirm that YOLOv8 and Mask R-CNN learn to detect reference TADs, new TADs and overlapping TADs (although absent in training data). All three show increased ChIP-Seq signal of the TAD-associated factors CTCF, RAD21 and H3K4me3 on their boundaries, emphasizing their biological reliability. Comparisons with reference TADs indicate that Mask R-CNN detects more TADs with less precise positioning, while YOLOv8 detects fewer TADs with lower nesting complexity but improved positioning.

**Keypoints:**

- Object detection methods are suited to detect TADs
- Object detection detects reasonable new TADs based on features learned from imperfect training data
- The main limitation and an important point for future work is the preciseness of bounding box positioning

## Introduction

Eukaryotic DNA, located in the nucleus, comprises of linear polymers of millions or even billions of nucleotides. The three-dimensional organization of DNA is not left to chance but highly determined by intramolecular contacts often facilitated by proteins (Rao et al., 2014). These contacts result in distinct structures that impact gene expression (Lieberman-Aiden et al., 2009). Topologically associated domains (TADs), structures of several 10 kbp to 1-2 Mbp, were identified as a core component of mammalian interphase chromatin (Dixon et al., 2012). Their formation is largely driven by cohesin-mediated loop extrusion, with CCCTC-binding factor (CTCF) acting as a directional insulator located at TAD boundaries (Fudenberg et al., 2016).

TADs serve as fundamental regulatory units (Lupiáñez et al., 2016) that contain regions of high chromatin interaction (Zhang et al., 2025), thereby facilitating promoter–enhancer communication within the same structural unit, while interactions across TAD boundaries are generally constrained (Fudenberg et al., 2016). Their functional relevance is further supported by observed conservation of TADs across different cell types and different species (Dixon et al., 2012; Fraser et al., 2015; Xu et al., 2024).

Since genome architecture is crucial for gene expression regulation, the knowledge of chromatin structure offers opportunities to better understand regulatory mechanisms. Adverse changes of TAD structures are linked to changes in promoter-enhancer interactions, resulting in an alteration of gene expression and can be associated with diseases such as cancer (Deng et al., 2022; Du et al., 2021; Giorgio et al., 2015; Lupiáñez et al., 2016; Valton & Dekker, 2016).

The structural composition of chromosomes can be examined using high-throughput chromosome conformation capture technology (Hi-C) (Lieberman-Aiden et al., 2009) quantifying the contact frequencies between genomic loci and thereby revealing the three-dimensional arrangement of chromatin. TADs appear as differently sized squares along the diagonal of Hi-C contact matrices (Zhang et al., 2025) and can be gapped, hierarchically nested and overlapping.

Each TAD is defined by two boundaries (genomic loci). Two TADs (TAD*_i,j_* and TAD*_m,n_*) with boundaries *i, j* and *m*, *n* have an inter-TAD gap (referred to as gapped TADs), if *i < j < m < n*. TAD*_m,n_* is nested within TAD*_i,j_* if *i ≤ m < n ≤ j* and *i ≠ m* xor *j ≠ n*. TAD*_i,j_* and TAD*_m,n_* overlap, if *i < m < j < n*. Figure S1 illustrates the types of interactions between pairs of TADs.

In Hi-C contact matrices, overlapping TADs are observed less frequently than nested TADs, which is likely because overlapping TADs result from bulk-sequencing, which captures an average view of the chromatin structure in a population of cells (Dali & Blanchette, 2017).

Although numerous TAD callers exist, they vary in their ability to identify distinct TAD interactions (Dali & Blanchette, 2017; Sefer, 2022; Zufferey et al., 2018). The Supplementary Methods provide a description of several tools and the types of interactions they capture. Many TAD callers are able to detect nested and/or gapped TADs (An et al., 2019; Rao et al., 2014; Weinreb & Raphael, 2015). However only a limited number of TAD callers (i.e. kerTAD (Maisuradze et al., 2024), Arrowhead (Rao et al., 2014), TADfit (Liu et al., 2022)) are able to identify overlapping, nested and gapped TADs.

Most TAD callers directly work with the Hi-C matrix or a graph representation (Xu et al., 2024). However, Hi-C matrices can be interpreted as images, which can be processed with computer vision methods like object detection. In object detection, object instances in an image are localized and classified (Pagire et al., 2024). These techniques have already been applied to detect corner dots of chromatin loops in Hi-C data (Chen et al., 2024) as well as micro domain boundaries in ChIA-Drop data (Tian et al., 2022).

In this work, we provide a proof of concept verifying that bounding-box based object-detection neural networks can successfully detect TAD structures in Hi-C matrices. For this purpose, we trained two models, YOLOv8 (Redmon et al., 2015) and Mask R-CNN (He et al., 2017), for the detection of TADs utilizing bounding boxes. A detailed explanation of both models is provided in the supplementary methods. To the best of our knowledge, no ground truth dataset exists and we therefore employed the OnTAD algorithm on Hi-C data to generate training data (which we refer to as reference TADs) since it has been shown to perform best in various categories of TAD-caller comparisons (Xu et al., 2024). Object detection techniques offer the advantages of high performance at very low runtime as well as the option for multiclass detection. To our knowledge, no existing method has yet employed object-detection techniques from computer vision - such as bounding-box-based detection or region-proposal-frameworks for the identification of TADs in Hi-C contact matrices.

YOLOv8 and Mask R-CNN are both faster than OnTAD and detected TADs show large agreement with OnTAD TADs. Moreover, both models detect overlapping TADs (which OnTAD cannot detect at all) and TADs that were not found by OnTAD. All three show peaks of CTCF, RAD21 and H3K4me3 ChIP-Seq signal on TAD boundaries. ChIP-Seq signals have not been fed to the networks and corroborate the validity of our predictions. While Mask R-CNN detects more TADs with less precise positioning (measured by ChIP-Seq peak height), YOLOv8 detects less TADs with accurate positioning.

## Materials and Methods

### Hi-C datasets and TADs

Two major problems in benchmarking and developing TAD detection algorithms are the unavailability of ground truth data and the limited resolution of Hi-C data. Potential methods to build datasets include: hand labeling of TADs, simulation of Hi-C data and TADs or usage of a (state-of-the-art) TAD caller. All approaches come with their own limitations. Hand labeling is very time-consuming and prone to small errors in the exact positioning of TAD boundaries, simulation induces biases in Hi-C data and TADs through preliminary assumptions over data properties, and TAD calling introduces an unknown number of false positives and false negatives. For our study we decided to train the models on real Hi-C data and TADs identified by a TAD caller, to observe how YOLOv8 and Mask R-CNN perform compared to a state-of-the-art method.

As shown by Xu et al. (2024) the TAD calling algorithm OnTAD performs best in different evaluation categories and they used it as their tool of choice for further evaluation of TAD hierarchy. This encouraged us to use OnTAD for the identification of TADs and comparison with the models we use.

We employ iteratively corrected (IC) (Imakaev et al., 2012) Hi-C data from the 4D Nucleome data portal (4DN) (Dekker et al., 2017; Reiff et al., 2022) in mcool format, focusing on datasets listed by the 4DN as "High-Resolution Hi-C Datasets" at 10 kbp resolution. IC implicitly removes technical and biological biases by iterative normalization of columns and rows of the Hi-C matrix.

The dataset used for training and validation is an in-situ Hi-C HepG2 (*hepatocellular carcinoma* cell line) dataset (4DNESC2DEQIJ) with 2.02 billion reads. The dataset used for testing is a micro-C (Goel et al., 2023) H1-hESC (embryonic stem cells) dataset (4DNES21D8SP8) with 3.22 billion reads after filtering. The fully disjunct test dataset enables us to evaluate the model performance on previously unseen data.

### Dataset processing

To produce data for training, validation and testing, we split chromosomes 1-22 of each dataset into a number of smaller Hi-C matrices. Each matrix contains 5.12 Mbp (512*512 pixels since we use a binsize of 10 kbp). The matrices overlap by half since we use a stepsize of 2.56 Mbp. This overlap can lead to data leakage, if overlapping matrices are split into train and validation dataset since they contain the same TADs. We consider this by splitting based on chromosomes. At the same time, the overlap ensures that we can detect all TADs up to the size of 2.56 Mbp, which lies above the upper limit of TAD sizes of approximately 2 Mbp (Dixon et al., 2012).

In iteratively corrected Hi-C data, the signal of diagonals decays as genomic distance between bins increases. This is known as distance-dependent decay (Lajoie et al., 2015). To reduce the discrepancy between main diagonal matrix values and off-diagonal matrix values, we preprocessed each Hi-C matrix using three transformations (C1, C2, C3) to generate a 3*512*512 pixel tensor as input for the models. Each of the transformations produces one of the three channels (Fig. 1 A, B, C). In the formulas below, *M* is the iteratively corrected matrix, *i* and *j* are row- and column-indices of a single matrix element. Addition of a pseudocount of 1e^-4^ is used to prevent log_10_(0) errors in the matrix. The notation *diag_|j-i|_* corresponds to the values of the *|j-i|*-th diagonal with *diag_0_* being the main diagonal. We use the absolute value of *j-i* since Hi-C matrices are symmetric.

**Figure 1:**
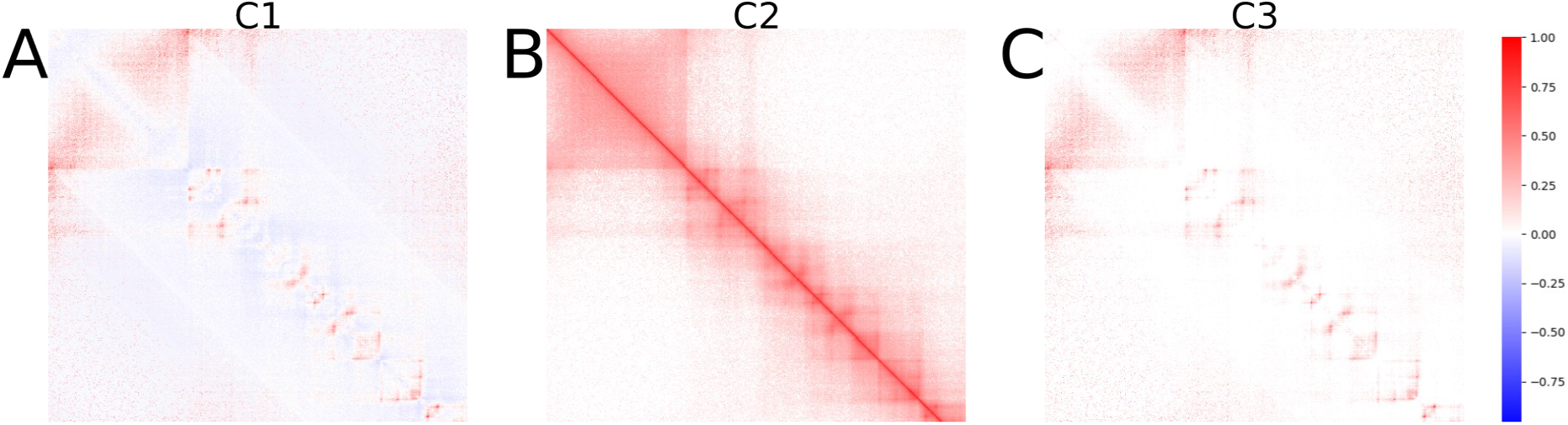
Display of a transformed Hi-C matrix used as three-channel model input. We process each 5.12 Mbp Hi-C submatrix in three ways and use the resulting matrices as model input. The matrix in panel A was processed by per-diagonal-z-score scaling, followed by max scaling (C1), the matrix in B was log transformed followed by min-max-scaling (C2) and the matrix in C was mean scaled, followed by filtering of negative values, max-scaling and the zeroing of the main diagonal as well as both one-off diagonals (C3).

In the first transformation (C1) we apply a per-diagonal-z-score scaling:

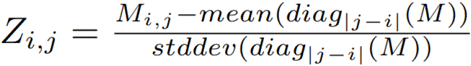

followed by max scaling based on the absolute values of the whole matrix *Z*:

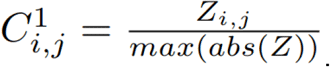

In the second transformation (C2) we upweight all non-main-diagonals by log transformation of the full matrix:

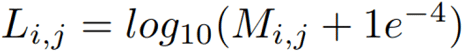

followed by min-max-scaling:

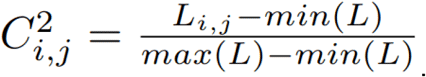

In the third transformation (C3) we set the diagonal means to zero:

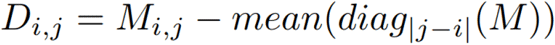

followed by removal of negative values:

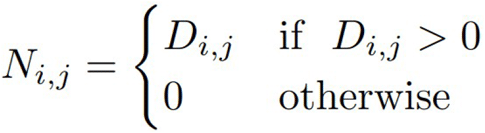

and max-scaling:

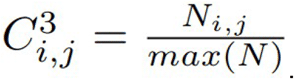

Afterwards, the main diagonal and both one-off diagonals of C3 are set to zero to reduce the signal emitted by self interacting loci.

After processing, the model input has the following properties:

- The values range from -1 to 1 (C1) and from 0 to 1 (C2, C3)
- Regions with high contact numbers like corner dots (which can be found in the upper right and lower left corner of some TADs in Hi-C matrices (Emerson et al., 2022)) and TAD boundaries are highlighted (Fig. 1A (C1) and C (C3))
- The square shape of TADs is highlighted (Fig. 1B (C2))
- Potential bias signal by the main diagonal and nearby diagonals is removed (Fig. 1C (C3))

We call TADs for both datasets using OnTAD with default parameters on iteratively corrected Hi-C matrices of single chromosomes. We then transform the coordinates of TADs on the chromosome to pixel coordinates in the images and remove all TADs that start or end on the image border because otherwise the high-contrast outline of the TAD is missing in the image.

### Model Training and Evaluation

YOLOv8 and Mask R-CNN were used on a local machine with a 13th Gen Intel Core i9-13950HX, 32 GB of RAM, and an NVIDIA RTX 3500 Ada. OnTAD was used on a SuperMicro SuperServer 4048B-TRFT with 4x Intel Xeon E7-8867v3 and 2048GB of RAM since the resources of the local machine were insufficient.

Chromosomes 1 and 3-22 of the HepG2 dataset are used for training; chromosome 2 is used for validation. The resulting HepG2 dataset consists of 1031 5.12 Mbp Hi-C matrices and 15283 unique reference TADs. 938 Hi-C matrices and 14030 TADs are used for training, 93 Hi-C matrices and 1253 TADs are used for validation. Chromosomes 1-22 of the H1-hESC dataset are used for testing. This dataset consists of 1041 5.12 Mbp Hi-C matrices and 20287 TADs.

The number of Hi-C matrices between the two datasets differs, since we do not use images without TADs. These images often contain large blank areas resulting from telo- or centromeric regions.

We trained YOLOv8 for 100 epochs and Mask R-CNN for 70 epochs due to early stopping. For each potential TAD the models return the bounding box coordinates and a confidence score. Since YOLOv8 and Mask R-CNN detections are not necessarily centered on the main diagonal, we average the upper left and lower right object instance coordinate: Each of the two coordinates is defined by two values *x* and *y*. A coordinate is on the main diagonal if *x=y*. If *x≠y* we solve this by setting *x=y=(x+y)/2*. From all TADs detected by YOLOv8 and Mask R-CNN, only TADs with a confidence score ≥ 0.5 are considered for further evaluation.

TADs detected by YOLOv8 and Mask R-CNN contain exact duplicates and highly overlapping TADs. This number increases further, since the submatrices we use overlap. To remove these duplicates, we use the intersection over union (IoU) of TAD pairs. IoU is commonly used in object detection tasks to determine the overlap between pairs of bounding boxes. It measures the ratio of intersection between two boxes with their union. Since TADs are centered on the main diagonal, we can define the IoU of two ascendingly sorted TADs with boundaries *i*, *j* and *m*, *n* as:

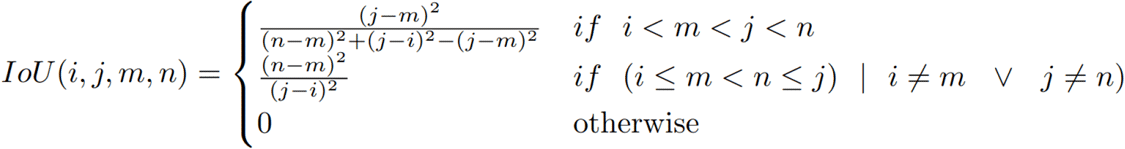

If two TADs detected by a model have an IoU above 0.5, we count them as duplicates and remove the TAD with the lower confidence score. This process is known as non-maximum suppression (NMS) (Felzenszwalb et al., 2010).

After confidence score and NMS filtering, we group the detected TADs of YOLOv8 and Mask R-CNN into nesting levels, similar to OnTAD. The nesting level of a TAD describes, how many TADs enclose it based on our definition of nested TADs (Fig. S1). If a TAD is not enclosed by others, its nesting level is 1.

We determine True Positives (TPs), False Positives (FPs), False Negatives (FNs) and calculate the True Positive Rate (TPR) and False Negative Rate (FNR) of the detected TADs. The True Negative Rate and False Positive Rate cannot be calculated since we have no True Negatives. A TP is defined as a TAD detected by YOLOv8 or Mask R-CNN that has an IoU ≥ 0.8 with a reference TAD. TPR and FNR are calculated as follows:

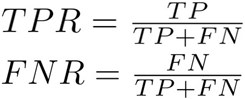

Since the TADs detected by each model contain overlaps, we calculate the number and sizes of overlaps and compare them between the two models.

### Biological Evaluation

To assess the biological relevance of detected TADs we compare the presence of TAD-associated factors (RAD21, CTCF and H3K4me3) at TAD boundaries between YOLOv8, Mask R-CNN and reference TADs based on ChIP-Seq data from ENCODE (ENCODE Project Consortium, 2012; Hitz et al., 2023; Kagda et al., 2025; Luo et al., 2020). These three factors are relevant for TAD formation, stability, transcriptional activity and thus provide biological support for correct TAD boundaries. RAD21 (HepG2: ENCFF242MRW, H1-hESC: ENCFF002NBT) is part of the cohesin complex through which DNA is extruded during loop extrusion, while DNA-bound CTCF (HepG2: ENCFF029RTI, H1-hESC: ENCFF332TNJ) interrupts the loop extrusion and thereby acts as an insulator (Kim et al., 2019). H3K4me3 (HepG2: ENCFF500VAH, H1-hESC: ENCFF760NUN) on the other hand is used as an epigenetic marker correlated with transcription and often found near promoters (Wang et al., 2023), which are partly regulated by TAD formation.

## Results

### Loss progression and model selection

The losses of both models decreased during training (Fig. S2). After training, we determined a single best epoch for each model and used the corresponding model to evaluate its performance on the previously unseen H1-hESC data. For Mask R-CNN we selected epoch 47 as the best run based on minimal losses of training and validation split (Fig. S2A, B). For YOLOv8, the selection of the best run is defined in the ultralytics code and based on a fitness score, which is a weighted sum of YOLOv8 performance metrics calculated on the validation data. The best fitness score was found at epoch 80, which does not coincide with an overfitting of YOLOv8 starting around epoch 85 (Fig. S2C, D).

### YOLOv8 and Mask R-CNN learn to detect reference TADs

Of the 20287 reference TADs in the test dataset, 11282 (=55.6%) and 8885 (=44.4%) were recovered by Mask R-CNN and YOLOv8, respectively (Fig. 2A, B). Although the IoUs of both models with reference TADs are high (Fig. S3A), the confusion matrices show large numbers of FPs (or OnTAD FNs) and FNs (or OnTAD FPs). This results in a True Positive Rate (TPR) of 0.556 and a False Negative Rate (FNR) of 0.444 for Mask R-CNN. The TPR of YOLOv8 is 0.438, the FNR is 0.562. Mask R-CNN detects 5356 TADs (=26.4%) more than OnTAD (Tab S1A). This results in a large number of FP TADs with varying overlap to OnTAD TADs. Two FPs with very low overlap to OnTAD are highlighted with yellow arrows in Fig. 2C. As YOLOv8 detects 5241 (=25.8%) TADs less than OnTAD (Tab S1A), most detections are FNs. Two FNs without a matching YOLOv8 TAD are highlighted with blue arrows in Fig. 2D.

**Figure 2:**
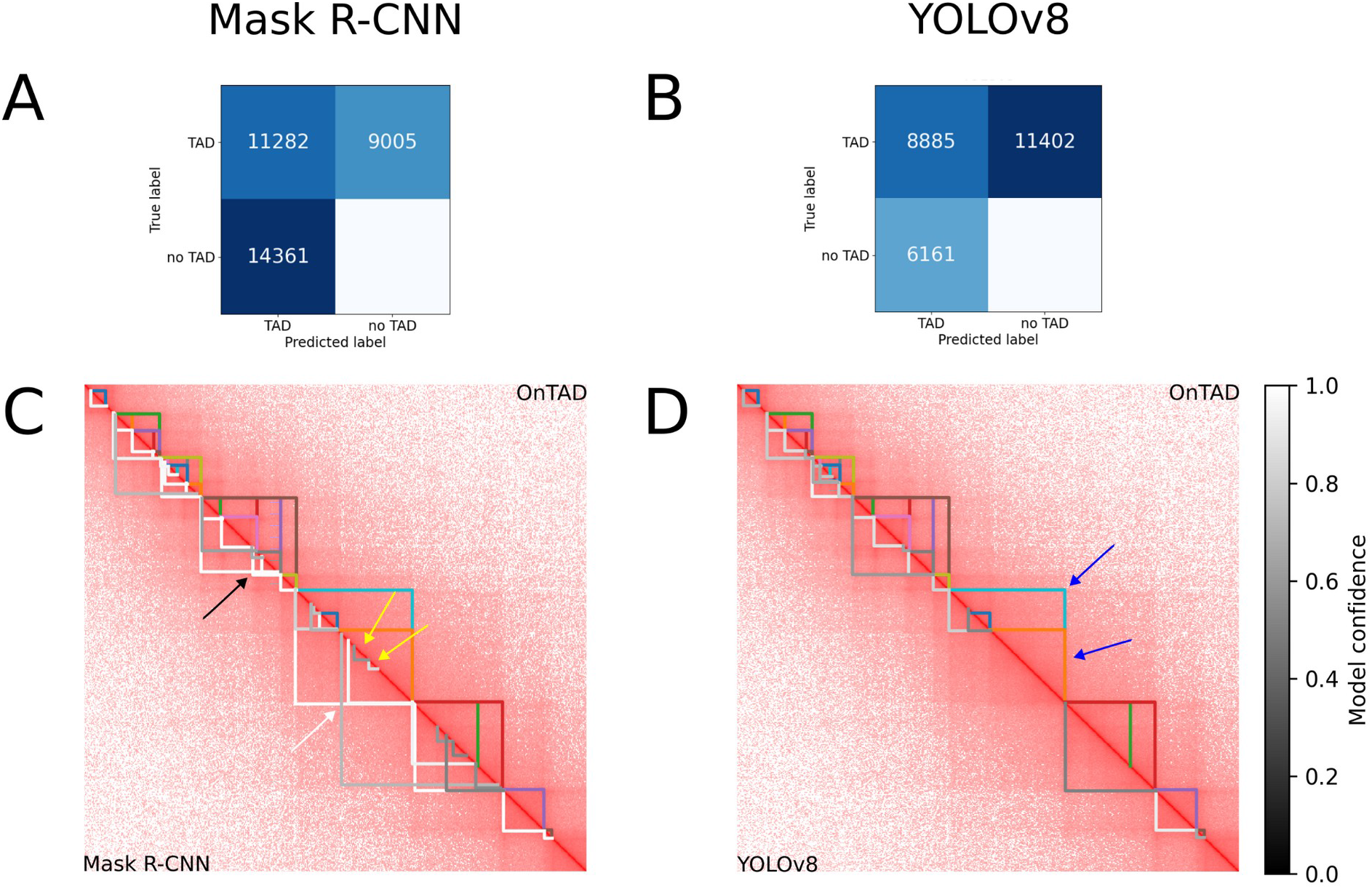
Overlap of TADs detected by YOLOv8 and Mask R-CNN with reference TADs. A and B show confusion matrices for Mask R-CNN- (A) and YOLOv8- (B) TADs detected in the H1-hESC dataset. A TAD is defined as TP if the maximum IoU with a reference TAD is ≥ 0.8. C and D show reference TADs in the upper triangular matrix and the predicted TADs of Mask R-CNN (C) and YOLOv8 (D) after confidence and NMS filtering in the lower triangular matrix. The iteratively corrected Hi-C matrix was log_10_ transformed followed by min-max scaling (C2) and shows the region chr18:25.600.000-30.720.000 bp of the H1-hESC dataset. The colors of reference TADs demarcate individual TADs, the colors of YOLOv8- and Mask R-CNN-TADs reflect the confidence score of the model (colorbar on the right). An example of a small (black arrow) and a big (white arrow) overlap of TADs is indicated in C. Two potential FPs of Mask R-CNN are highlighted with yellow arrows, two FNs of YOLOv8 are highlighted with blue arrows.

The Mask R-CNN TADs displayed in Fig. 2C show an additional property missing in reference TADs: some of the displayed TADs (highlighted with black and white arrows) overlap. YOLOv8 TADs do not show overlaps in the displayed region, although the model is able to place overlapping bounding boxes.

### TAD boundaries found by YOLOv8 and Mask R-CNN show concordance with biological data

To assess whether the detected TAD boundaries align with biological markers we calculated the average fold-change of CTCF, RAD21 and H3K4me3 ChIP-Seq data around all unique start and stop boundaries of detected TADs (Fig. 3). We then compare the average fold-change between models.

**Figure 3:**
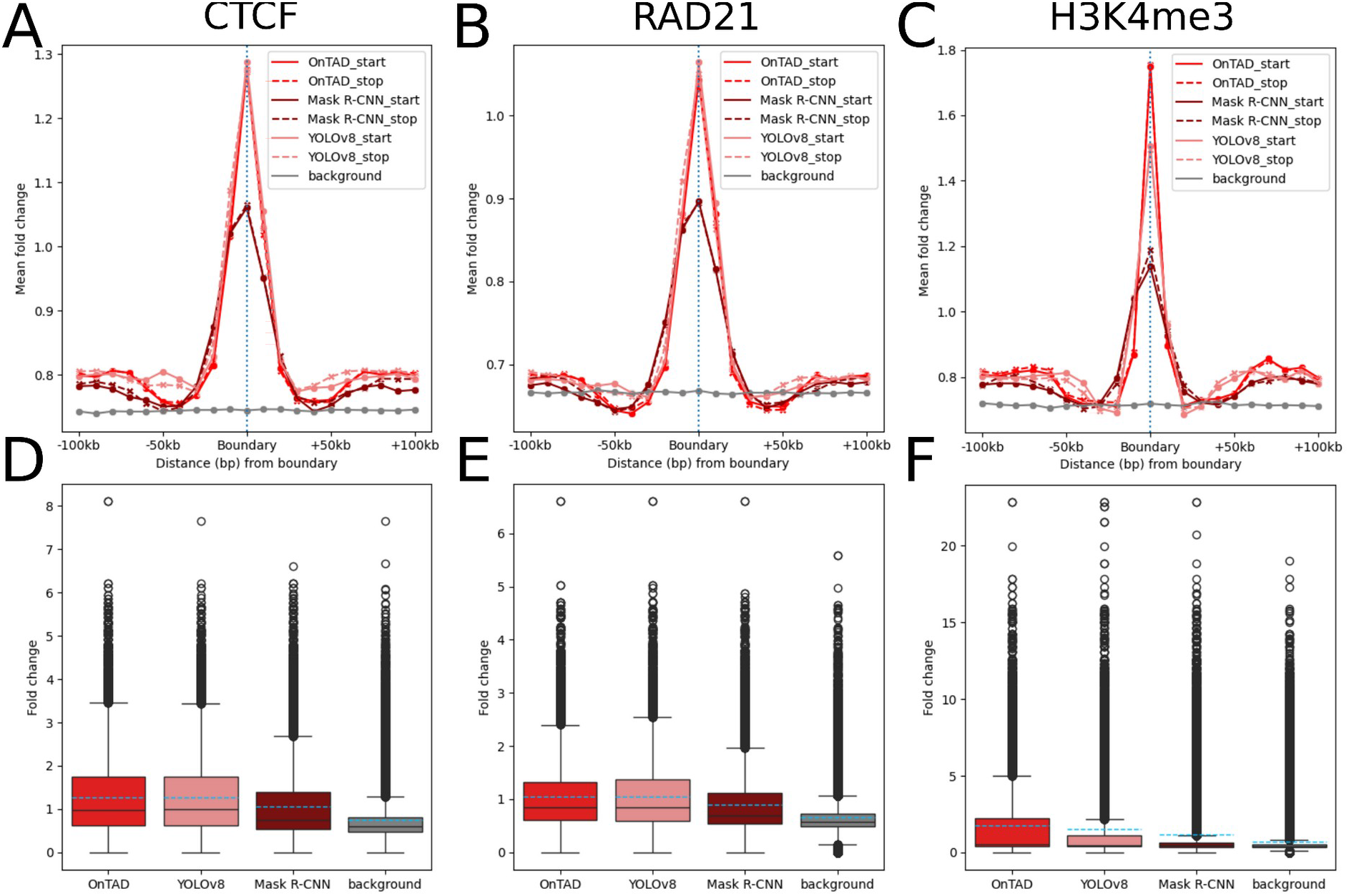
ChIP-Seq signal of CTCF (A, D), RAD21 (B, E) and H3K4me3 (C, F) at TAD boundaries (excluding duplicates) of reference TADs, TADs predicted by YOLOv8 and Mask R-CNN on the H1-hESC data used for testing. In A, B and C, the averaged fold-change in a 200 kbp window around the boundary (21 bins) is shown to display differences in ChIP-Seq signal. The background signal of randomly drawn positions is added in grey. Blacklist regions (Amemiya et al., 2019) are excluded from background calculation. The position of the predicted boundary is shown as a blue dotted line. In D, E and F, boxplots show the ChIP-Seq signal of boundaries (excluding duplicates). Dashed lines mark the arithmetic means, solid lines the medians. Grey boxes show the background signal of randomly drawn positions.

The TADs detected by both models and the reference show an increase of ChIP-Seq signal on the boundaries of found TADs. YOLOv8 matches the reference CTCF and RAD21 signal on unique TAD boundaries, while Mask R-CNN shows lower fold-changes for both factors (Tab S1D). The H3K4me3 signal is the highest for reference boundaries, followed by YOLOv8 and Mask R-CNN.

Since both models detect many FPs, we split detections into the fraction of TADs with a reference IoU < 0.2 (*new*), the fraction of TPs with a reference IoU ≥ 0.8 (*present*) and the remaining TADs (*intermediate*) (Fig. S3B). YOLOv8 detects 1597 *new* TADs (=10.6% of detections), Mask R-CNN 4070 (=15.9% of detections).

To examine whether the *new* TADs are FPs lacking biological support or reasonable TADs that are missing in the reference, we compared the ChIP-Seq signal at boundaries between the groups *new* and *present*. We found that all groups show an increase of ChIP-Seq signal on their boundaries compared to the surrounding region, but the peak heights differ (Fig. 4). The ChIP-Seq signal at boundaries of YOLOv8 TADs is generally higher than that of Mask R-CNN boundaries. The ChIP-Seq signal on boundaries of *present* and *new* differs more for Mask R-CNN TADs. In addition, the peak shape of Mask R-CNN boundaries is less sharp compared to YOLOv8, especially for CTCF and RAD21.

**Figure 4:**
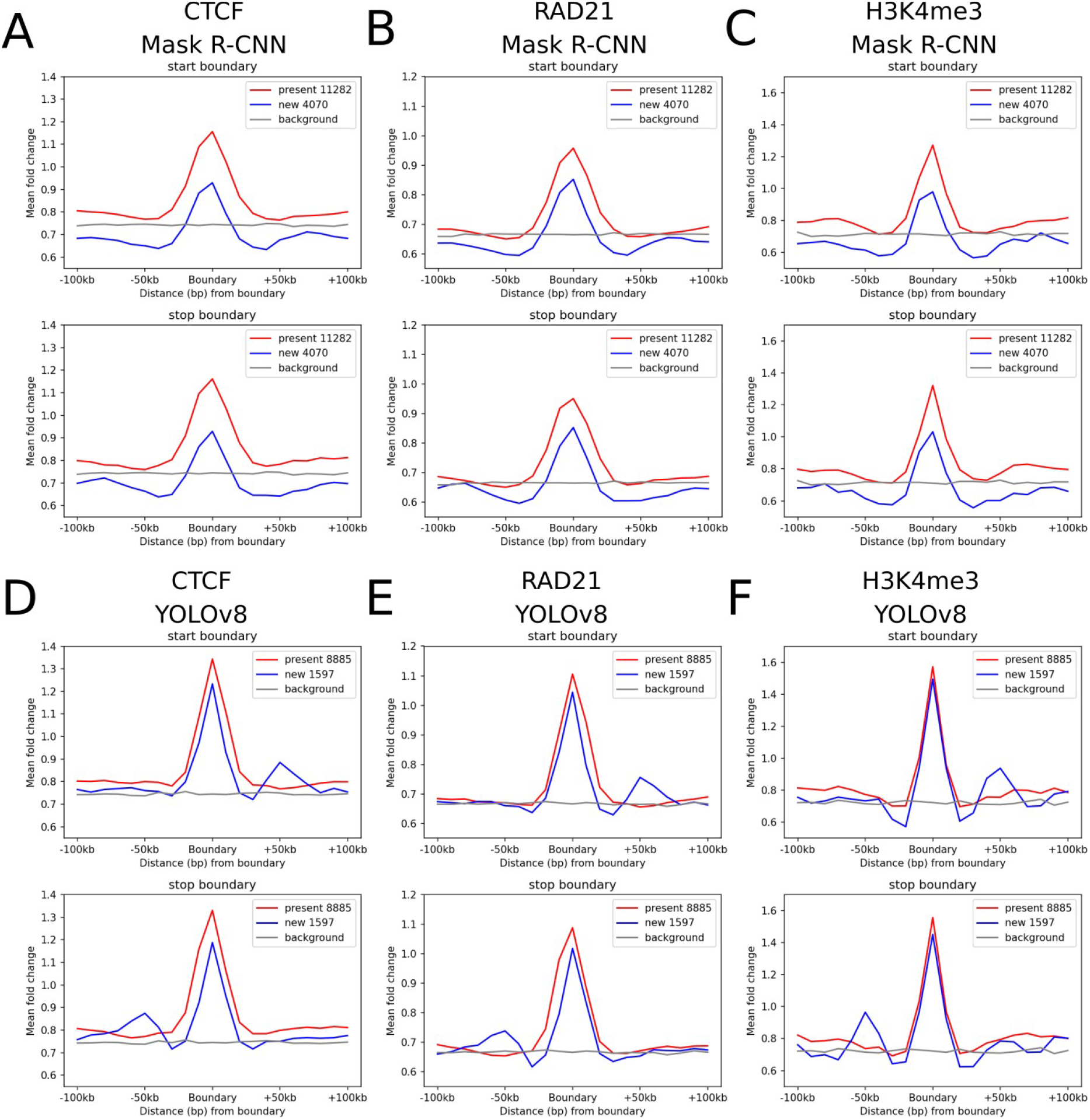
Comparison of ChIP-Seq signal at boundaries of *new* TADs (blue) compared to *present* TADs (red) detected in the test dataset. ChIP-Seq fold change signals for CTCF (A, D), RAD21 (B, E) and H3K4me3 (C, F) were averaged over *new* TADs (blue) as well as *present* TADs (red) for Mask R-CNN (A-C) and YOLOv8 (D-F). The boundaries are separated into TAD-start and -stop. Grey lines show the background signal. Blacklist regions (Amemiya et al., 2019) are excluded from background calculation.

The separation of boundaries into start and stop shows an additional trend: In Figure 4 D, E and F the boundaries of *new* TADs detected by YOLOv8 show a second increase in ChIP-Seq signal approximately 50 kbp upstream of the stop boundary and downstream of the start boundary. This peak is present in CTCF, RAD21 and H3K4me3. *New* TADs detected by Mask R-CNN show only a minor increase upstream of the stop boundary and downstream of the start boundary.

### The unrestricted object detection of YOLOv8 and Mask R-CNN facilitates the detection of overlapping TADs

One advantage of YOLOv8 and Mask R-CNN compared to most TAD callers is their ability to detect objects without restrictions in object positioning. OnTAD for example cannot detect overlapping TADs and therefore, the reference data we use for training contains no overlapping TADs. In contrast, Fig. 2A shows that Mask R-CNN TADs do overlap, spanning a variety of overlap sizes. Based on this finding, we counted the number of overlapping TADs and their overlap sizes in YOLOv8 and Mask R-CNN detections (Fig. 5). In the test dataset 20761 (=81.0%) of 25643 TADs detected by Mask R-CNN overlap with at least one other TAD. The 20761 TADs build 25308 overlaps. In contrast, only 4706 (=31.3%) of 15046 TADs detected by YOLOv8 overlap and build 3220 overlaps (Tab S1F).

**Figure 5:**
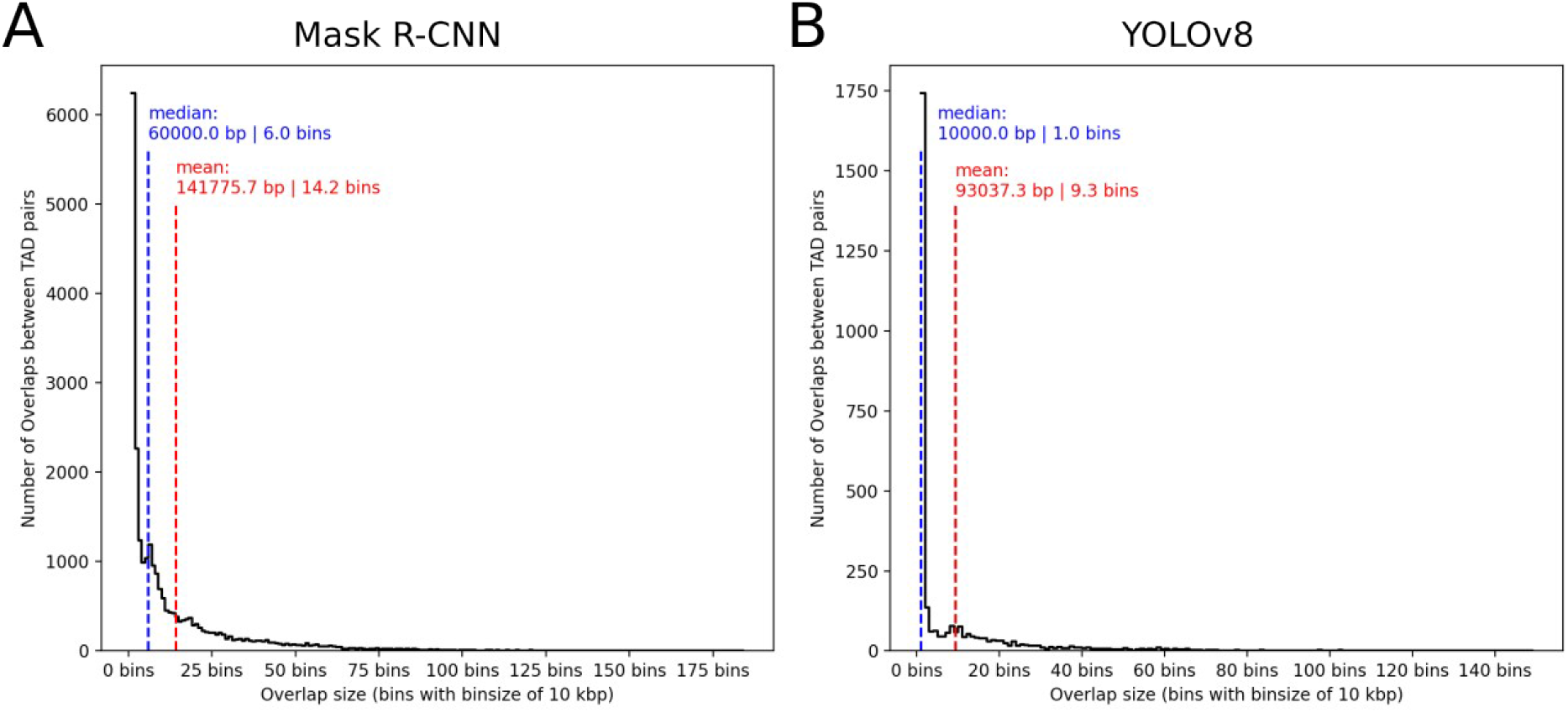
Overlap sizes of overlapping TAD-pairs detected by Mask R-CNN (A) and YOLOv8 (B) in the test dataset. The plots include all overlaps between TAD pairs and overlap sizes are shown in bins. Mean and median are added as vertical lines.

We noticed that most of the overlaps are small and comprise only a few bins (Fig. 5). This holds especially for YOLOv8, where the median size of overlaps is a single bin. 7559 overlaps (=29.9%) of Mask R-CNN TADs and 935 overlaps (=29.0%) of YOLOv8 TADs are larger than the corresponding mean overlap sizes of 14.2 bins and 9.3 bins.

### YOLOv8 and Mask R-CNN show faster runtimes compared to the reference

During model training, Mask R-CNN took 167.8 ± 1.3 seconds per epoch (one epoch includes training on chr1, chr3-22 and validation on chr2) while YOLOv8 took 29.9 ± 1.3 seconds for a single epoch (Fig. S4). For a runtime comparison with OnTAD, we repeated TAD detection on chromosome 1-22 of the H1-hESC dataset 10 times with OnTAD, YOLOv8 and Mask R-CNN. While OnTAD took 1089.5 ± 417.3 seconds, Mask R-CNN took 68.1 ± 5.4 seconds and YOLOv8 took 22.5 ± 1.3 seconds. Leveraging GPU parallelism, Mask R-CNN is 16-fold and YOLOv8 is 50-fold and faster than OnTAD, which is only available for CPU compute, respectively.

## Discussion

After training, YOLOv8 and Mask R-CNN recover many TPs although with large numbers of supposedly FPs and FNs. These observations are tied to the chosen confidence score and NMS filtering, which are discussed in the supplementary section “Selection of data augmentation and thresholds”. Our relatively low NMS of 0.5 may for example remove some non-duplicate TADs from our results.

Analysis of ChIP-Seq signal shows that both models and OnTAD detect biologically reliable TADs reflected by an increase of ChIP-Seq signal on their boundaries. While YOLOv8 reaches the reference in terms of CTCF and RAD21 sginal at boundaries in the training and test dataset, the signal of Mask R-CNN boundaries is always lower. On the one hand, this indicates that YOLOv8 places the bounding boxes more precisely than Mask R-CNN. This observation is supported by the difference between one-stage (YOLOv8) and two-stage (Mask R-CNN) detection processes: In the first stage of Mask R-CNN, the region proposal network (RPN) detects a number of region proposals, each defined by a bounding box. If a bounding box would match a TAD exactly, the resulting region proposal would crop the difference in contrast at the TAD boundary which complicates the boundary detection. On the other hand, YOLOv8 detects fewer TADs than Mask R-CNN and most of them have a low nesting level (Fig. S7, S9). This may contribute to the increase in CTCF ChIP-Seq signal on TAD boundaries compared to Mask R-CNN, since larger TADs with a lower nesting level show increased CTCF signal on their boundaries compared to small TADs with a high nesting level (An et al., 2019). However, the performance of YOLOv8 and Mask R-CNN is tied to the reference training data generated with OnTAD. Since we do not have a real reference dataset, the quality of the training dataset is unknown and the set of TADs is most likely incomplete.

This assumption is supported by the TADs with very small reference overlap (*new*, originally FPs) which show an increase in ChIP-Seq singal on their boundaries. It confirms that the models do not simply learn to imitate the reference algorithm but instead learn TAD-specific features present in the Hi-C matrices and, based on these features, detect new TADs.

For YOLOv8 the validity of *new* TADs is further supported by the peaks upstream and downstream of the boundaries (Fig. 4D-F). These peaks are located at a distance of approximately 50 kbp, which matches the predominant size of the *new* TADs (Fig. S5). Theses second peaks are the start- and stop-boundaries corresponding to the centered boundary in Figure 4. Since *new* TADs detected by Mask R-CNN have more variety in size, the peaks are only visible as a minor increase in ChIP-Seq signal.

Although the training data contained no overlapping TADs, we found a variety of overlaps in TADs detected by YOLOv8 and Mask R-CNN. On the one hand, the larger overlaps confirm that the unrestricted object detection reveals a group of TADs missing in the training data, which exhibit enrichment of CTCF, RAD21 and H3K4me3 (Fig. S6). On the other hand, the large number of tiny overlaps, especially in Mask R-CNN detections, highlights a limitation of the models: Since overlaps of e.g. a single bin are not described in literature, we hypothesize that both models struggle to place bounding box boundaries with high precision. This is further supported by the high number of unique boundaries compared to detected TADs (Tab. S1, Tab. S2).

In terms of runtime, we observe a major improvement compared to the reference. This is expected, as both models were originally designed for fast detection to be applicable in tasks like autonomous driving. The speed difference between YOLOv8 and Mask R-CNN is rooted in the difference between models. Mask R-CNN consists of two neural networks which detect successively while YOLOv8 detects in a single step. We have to note that the runtime comparability of the reference and the two models is limited since they rely on different hardware components.

## Conclusion

In this study, we examined the applicability of the object detection models YOLOv8 and Mask R-CNN in TAD detection.

We built two datasets from Hi-C matrices to train and evaluate the models by applying three transformations to overlapping regions of the Hi-C matrix to highlight different properties of the data. The TADs for both datasets were generated by using the TAD calling algorithm OnTAD, which we chose based on a review of TAD callers.

YOLOv8 and Mask R-CNN detect biologically reliable TP TADs but struggle with exact TAD positioning and high numbers of FP and FN detections. We found that the FP TADs contain a fraction of new TADs which show ChIP-Seq signal peaks of TAD-associated factors on their boundaries. This confirms that the models are no perfect out-of-the-box TAD detectors but have the potential to detect biologically reliable TADs not found by a reference method. Future work should address the precise detection of TAD boundaries, measuring dataset quality, the impact of dataset size and transfer learning on detection quality, an extension to multiclass detections (for example chromatin loops) and a combination of complementary input data as recently proposed by Dekker et al. (2025).

## Funding

This work was supported by the Forschungscampus Mittelhessen Project 2026_1_02.

## Data availability

We used only publicly available datasets from 4Dnucleome and ENCODE in this study. All generated data and all of the publicly available data (except Hi-C) is available at https://doi.org/10.5281/zenodo.23389677. The Hi-C data can be downloaded from the 4Dnucleome.

## Code availability

Code for model training, evaluation and visualization is available at Zenodo https://doi.org/10.5281/zenodo.23389677

## Supplementary data

### Supplementary Tables

**Table S1:** Numbers of reference TADs, TADs detected by YOLOv8 and Mask R-CNN on the H1-hESC dataset used for testing. A shows the number of TADs detected by all methods and the number of TADs grouped into nesting levels (TADs with a nesting level ≥ 4 are summed in the “4+” group). TADs of YOLOv8 and Mask R-CNN include all NMS filtered TADs with a confidence score ≥ 0.5. B shows the number of TAD boundaries, unique TAD boundaries and unique TAD boundaries grouped into nesting levels. If a single TAD boundary is shared between multiple TADs, the nesting level of the boundary is the highest nesting level of the TADs. C shows the mean sizes of TADs grouped into nesting levels. D shows the mean and median ChIP-Seq fold change values computed over all unique TAD boundaries detected by a method. The ChIP-Seq factors include CTCF, RAD21 and H3K4me3. E shows the number of TADs split into the groups *present* (IoU >= 0.8), *intermediate* (0.8 > IoU ≥ 0.2) and *new* (IoU < 0.2) based on their maximum IoU with reference TADs. F shows the number of TADs that overlap with one or more TADs detected by the same method, the number of overlaps as well as the mean and median sizes of these overlaps.

|  |  | OnTAD | YOLOv8 | Mask R-CNN |
| --- | --- | --- | --- | --- |
| **A** | Total TADs | 15284 | 9286 | 21106 |
|  | Level 1 | 4070 | 5008 | 6598 |
|  | Level 2 | 4869 | 3292 | 6174 |
|  | Level 3 | 3480 | 868 | 4439 |
|  | Level 4+ | 2865 | 118 | 3895 |
| **B** | Total Boundaries | 30568 | 18572 | 42212 |
|  | Unique Boundaries | 11705 | 12199 | 30882 |
|  | Level 1 | 1385 | 5565 | 8301 |
|  | Level 2 | 3214 | 4914 | 8943 |
|  | Level 3 | 3452 | 1501 | 6998 |
|  | Level 4+ | 3654 | 219 | 6640 |
| **C** | Mean size Level 1 | 5.627 | 5.462 | 5.575 |
|  | Mean size Level 2 | 5.393 | 5.275 | 5.377 |
|  | Mean size Level 3 | 5.315 | 5.158 | 5.269 |
|  | Mean size Level 4+ | 5.249 | 5.095 | 5.169 |
| **D** | Mean CTCF FC on boundary | 0.927 | 0.935 | 0.807 |
|  | Median CTCF FC on boundary | 0.747 | 0.751 | 0.659 |
|  | Mean RAD21 FC on boundary | 0.937 | 0.949 | 0.818 |
|  | Median RAD21 FC on boundary | 0.769 | 0.771 | 0.678 |
|  | Mean H3K4me3 FC on boundary | 1.177 | 1.146 | 0.931 |
|  | Median H3K4me3 FC on boundary | 0.683 | 0.667 | 0.651 |
| **E** | *Present* | 15284 | 5737 | 8034 |
|  | *Intermediate* | 0 | 2846 | 8611 |
|  | *New* | 0 | 703 | 4461 |
| **F** | Overlapping TADs | 0 | 2494 | 17122 |
|  | Number of Overlaps | 0 | 1631 | 20322 |
|  | Mean Overlap size (bins) | 0.0 | 10.1 | 13.6 |
|  | Median overlap size (bins) | 0.0 | 1.0 | 6.0 |

**Table S2:** Numbers of reference TADs, TADs detected by YOLOv8 and Mask R-CNN on the HepG2 dataset used for training. A shows the number of TADs detected by all methods and the number of TADs grouped into nesting levels (TADs with a nesting level ≥ 4 are summed in the “4+” group). TADs of YOLOv8 and Mask R-CNN include all NMS filtered TADs with a confidence score ≥ 0.5. B shows the number of TAD boundaries, unique TAD boundaries and unique TAD boundaries grouped into nesting levels. If a single TAD boundary is shared between multiple TADs, the nesting level of the boundary is the highest nesting level of the TADs. C shows the mean sizes of TADs grouped into nesting levels. D shows the mean and median ChIP-Seq fold change values computed over all unique TAD boundaries detected by a method. The ChIP-Seq factors include CTCF, RAD21 and H3K4me3. E shows the number of TADs split into the groups *present* (IoU >= 0.8), *intermediate* (0.8 > IoU ≥ 0.2) and *new* (IoU < 0.2) based on their maximum IoU with reference TADs. F shows the number of TADs that overlap with one or more TADs detected by the same method, the number of overlaps as well as the mean and median sizes of these overlaps.

|  | OnTAD | YOLOv8 | Mask R-CNN |
| --- | --- | --- | --- |
| **A** Total TADs | 20287 | 15046 | 25643 |
| Level 1 | 4102 | 6594 | 6845 |
| Level 2 | 6131 | 6047 | 6946 |
| Level 3 | 5139 | 2018 | 5982 |
| Level 4+ | 4915 | 387 | 5870 |
| **B** Total Boundaries | 40574 | 30092 | 51286 |
| Unique Boundaries | 14385 | 18012 | 36181 |
| Level 1 | 571 | 5852 | 8040 |
| Level 2 | 3022 | 8203 | 9193 |
| Level 3 | 4654 | 3261 | 9041 |
| Level 4+ | 6138 | 696 | 9907 |
| **C** Mean size Level 1 | 5.689 | 5.449 | 5.602 |
| Mean size Level 2 | 5.417 | 5.262 | 5.390 |
| Mean size Level 3 | 5.332 | 5.162 | 5.267 |
| Mean size Level 4+ | 5.273 | 5.108 | 5.161 |
| **D** Mean CTCF FC on boundary | 1.277 | 1.280 | 1.063 |
| Median CTCF FC on boundary | 0.986 | 1.005 | 0.756 |
| Mean RAD21 FC on boundary | 1.044 | 1.057 | 0.896 |
| Median RAD21 FC on boundary | 0.852 | 0.850 | 0.688 |
| Mean H3K4me3 FC on boundary | 1.752 | 1.501 | 1.163 |
| Median H3K4me3 FC on boundary | 0.504 | 0.474 | 0.446 |
| **E** *Present* | 20287 | 8885 | 11282 |
| *Intermediate* | 0 | 4564 | 10291 |
| *New* | 0 | 1597 | 4070 |
| **F** Overlapping TADs | 0 | 4706 | 20761 |
| Number of Overlaps | 0 | 3220 | 25308 |
| Mean Overlap size (bins) | 0.0 | 9.3 | 14.2 |
| Median overlap size (bins) | 0.0 | 1.0 | 6.0 |

### Supplementary Figures

**Figure S1:**
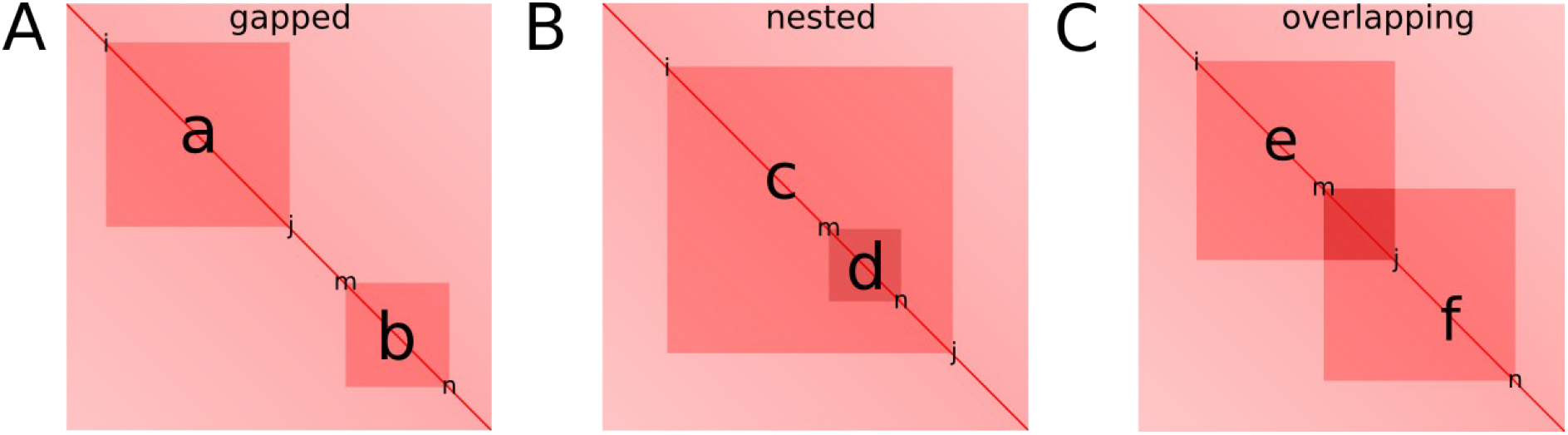
Schematic illustration of interaction types between TADs in a Hi-C matrix. TADs are displayed as red squares centered on the main diagonal and named with single lowercase letters. The possible types of interactions are gapped TADs (A), nested TADs (B) and overlapping TADs (C). TAD c has nesting level of 1 since no other TAD completely encloses it, while TAD d has nesting level of 2.

**Figure S2:**
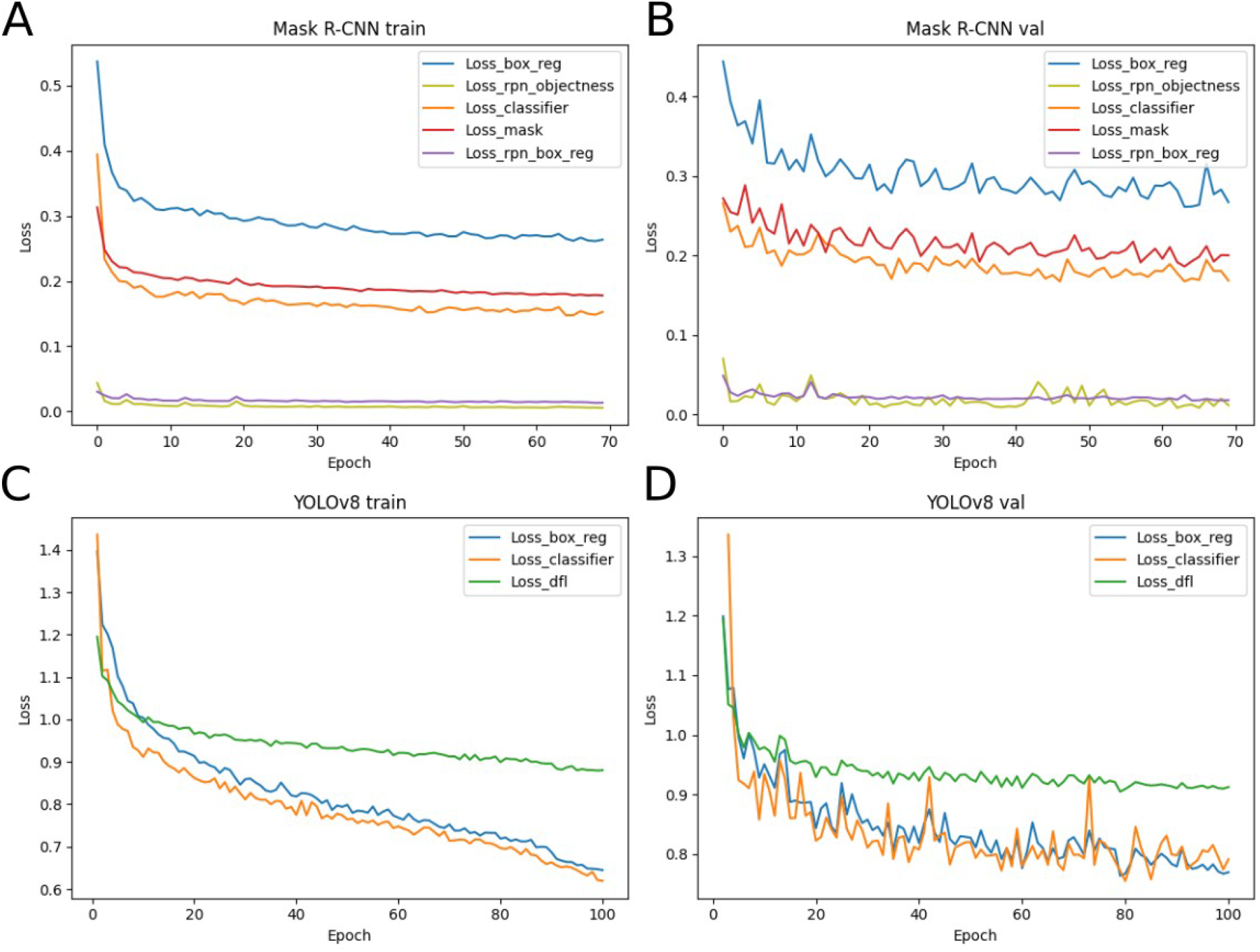
Model losses on training and validation data. Losses of Mask R-CNN (A, B) and YOLOv8 (C, D) during training (A, C) and validation (B, D). Colors correspond to different losses.

**Figure S3:**
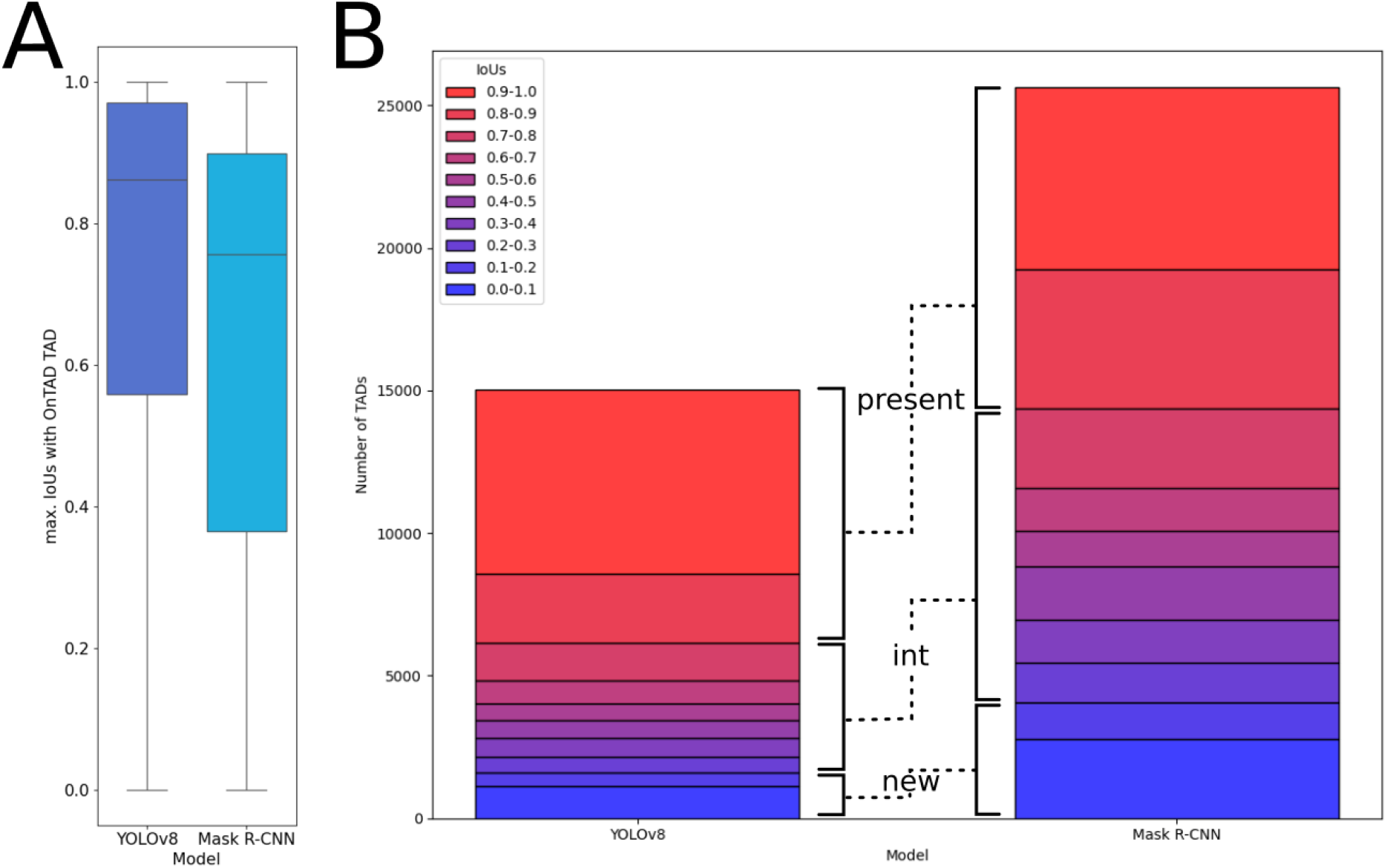
Overlap of TADs detected by YOLOv8 and Mask R-CNN with reference TADs in the test dataset. A shows boxplots of the maximum IoUs of all YOLOv8 and Mask R-CNN TADs with reference TADs. B shows the IoU-based grouping of TADs detected by both models with reference TADs. For each detected TAD of both models, the maximum IoU with a reference TAD is calculated. This IoU is used to assign the TAD to *present* in the reference TADs, *intermediate* or *new*. The black brackets mark the corresponding intervals in which a TAD is labeled as *present* (1.0 ≥ IoU ≥ 0.8) or *new* (0.2 < IoU ≤ 0.0). All TADs with 0.8 > IoU ≥ 0.2 are labeled as *intermediate* and excluded from further analysis since we cannot definitely add them to one of the two groups.

**Figure S4:**
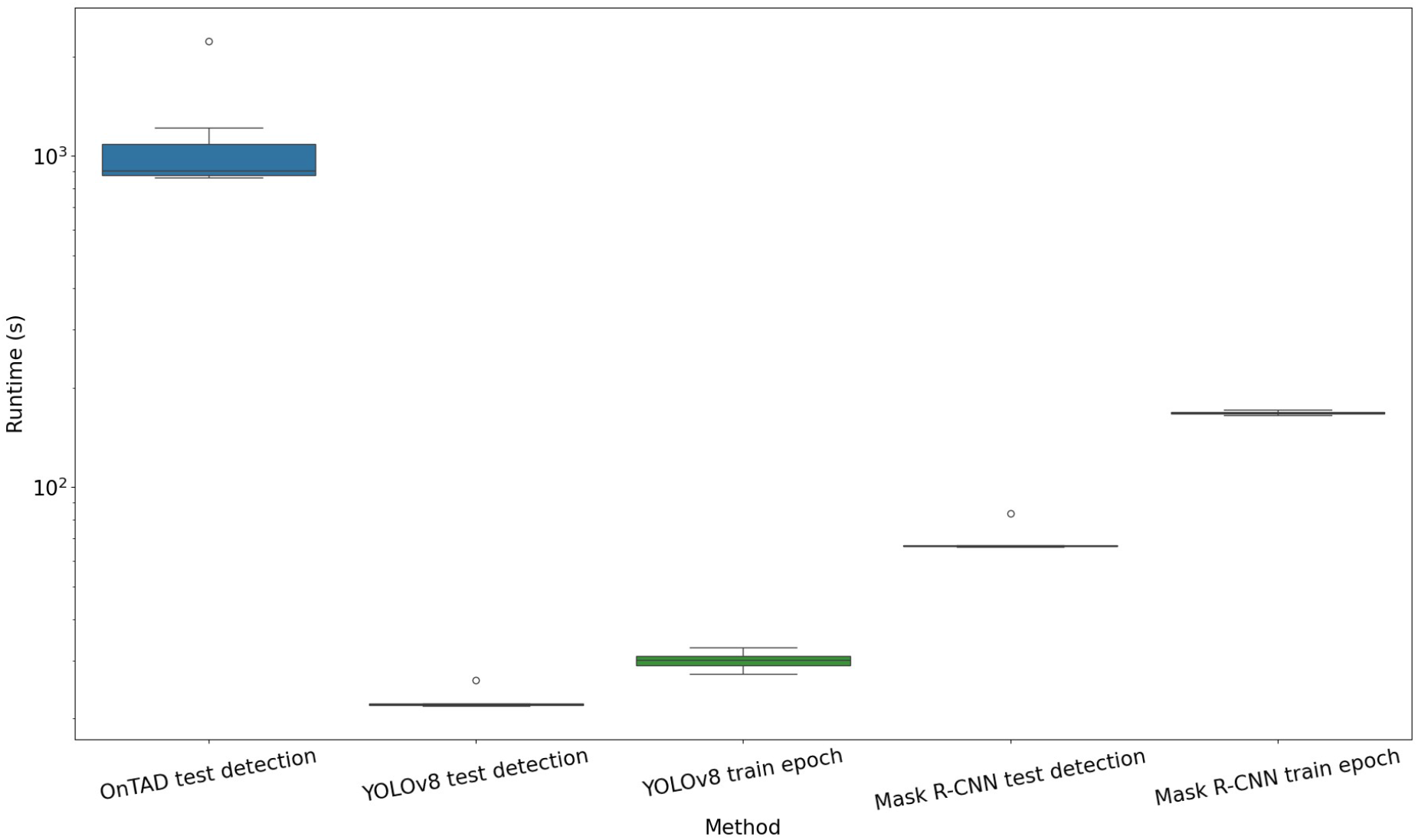
Runtime Comparison of OnTAD, YOLOv8 and Mask R-CNN. “test detection” refers to detection of TADs in chromosomes 1-22 of the H1-hESC dataset, which was repeated 10 times. “train epoch” shows the runtime of a single epoch during training of YOLOv8/Mask R-CNN on chromosomes 1 and 3-22 as well as validation on chromosome 2 of the HepG2 dataset.

**Figure S5:**
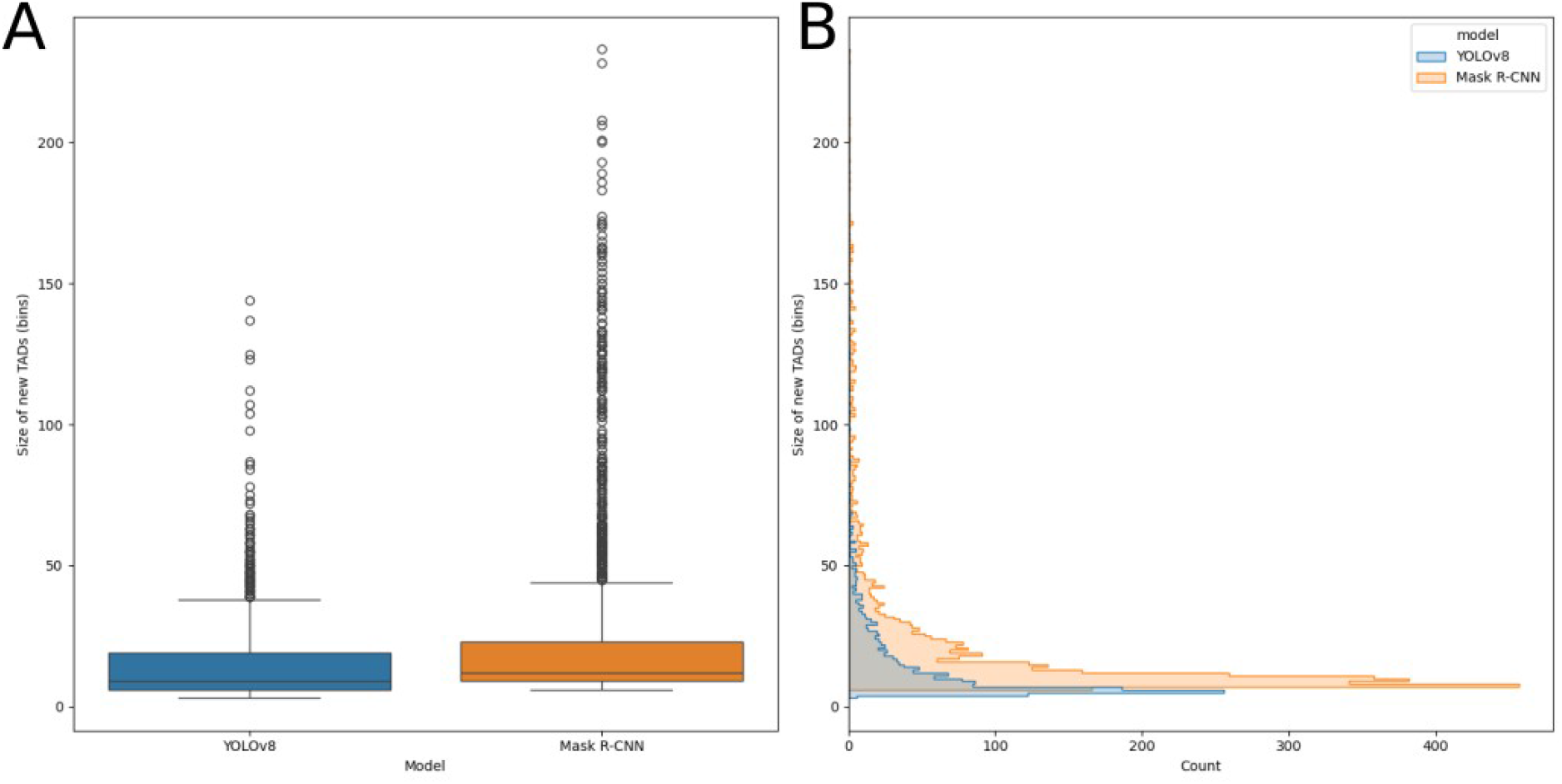
Sizes of *new* TADs predicted by YOLOv8 and Mask R-CNN. A: Boxplot of TAD-sizes for each model in bins. B: Histogram of TAD-sizes for each model in bins.

**Figure S6:**
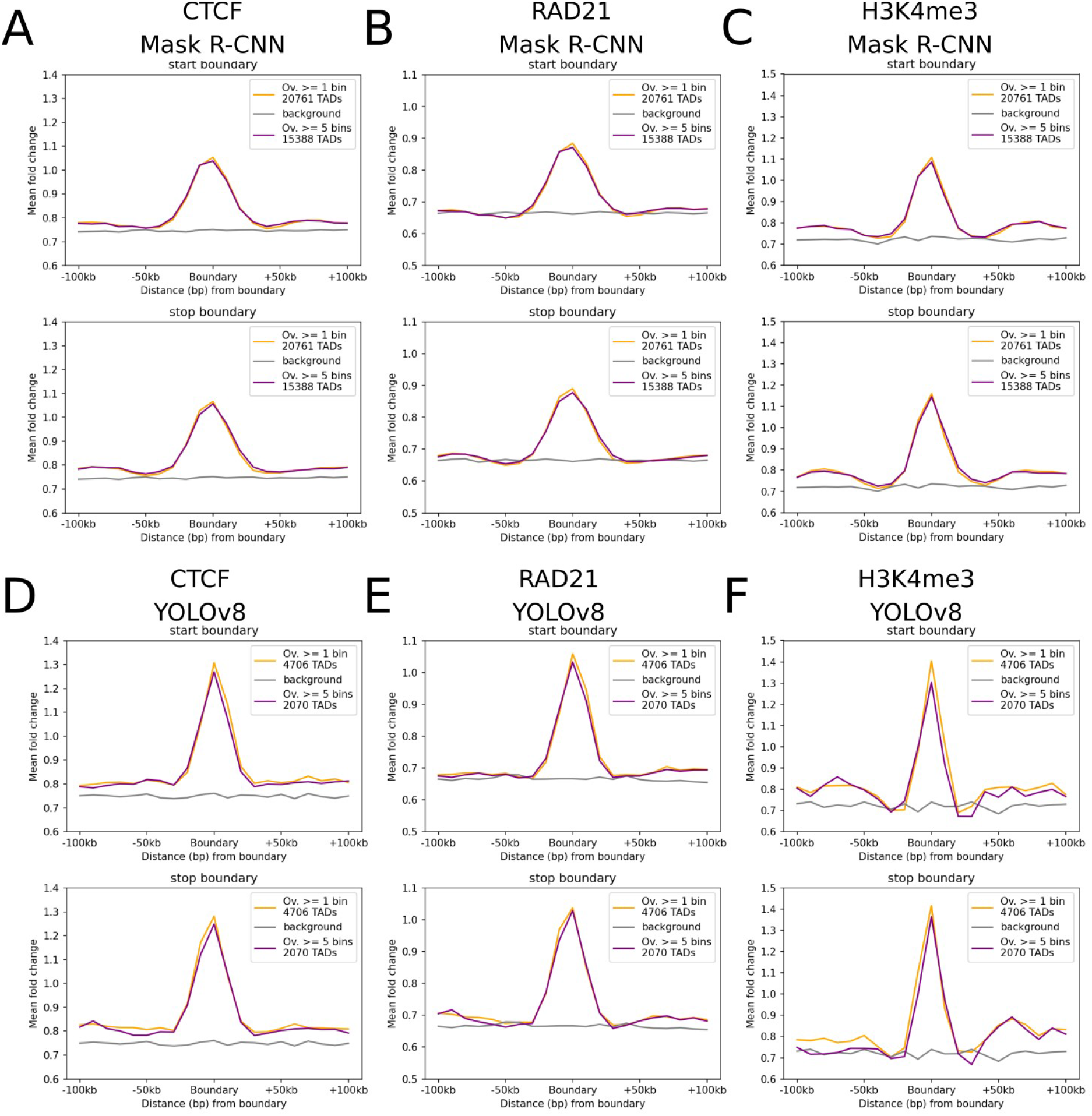
Comparison of ChIP-Seq signal at boundaries of overlapping TADs for two different minimum overlap sizes. ChIP-Seq fold change signals for CTCF (A, D), RAD21 (B, E) and H3K4me3 (C, F) were averaged over two groups of overlapping TADs for Mask R-CNN (A-C) and YOLOv8 (D-F). The purple lines include all overlaps, the yellow lines include all overlaps ≥ 5 bins. Grey lines show the background signal. Blacklist regions (Amemiya et al., 2019) are excluded from background calculation. The boundaries are separated into TAD-start and -stop.

**Figure S7:**
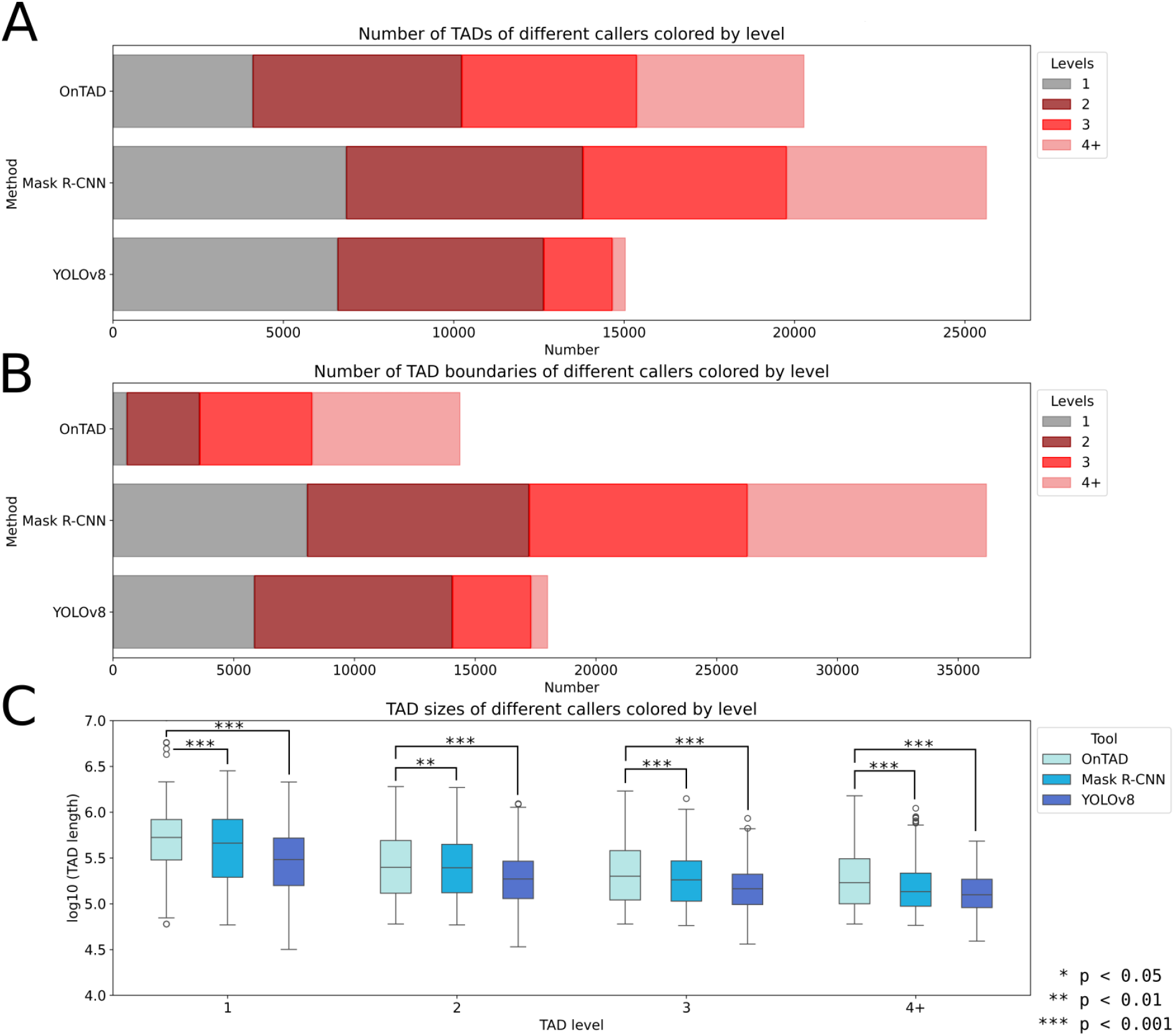
Characteristics of reference TADs, TADs detected by Mask R-CNN and YOLOv8 in the test dataset. TADs are subdivided according to their nesting level (TADs with a nesting level ≥ 4 are summed in the “4+” group). A shows the number of detected TADs. B shows the number of unique boundaries of TADs subdivided based on TAD level: If one genomic position is the boundary of multiple TADs, we select the highest level of the TADs for B. C shows the sizes of TADs. Significance levels of differences in TAD sizes between reference and both models were calculated using the two-sided Mann-Whitney U test and p-values were adjusted for multiple testing by Benjamini-Hochberg.

**Figure S8:**
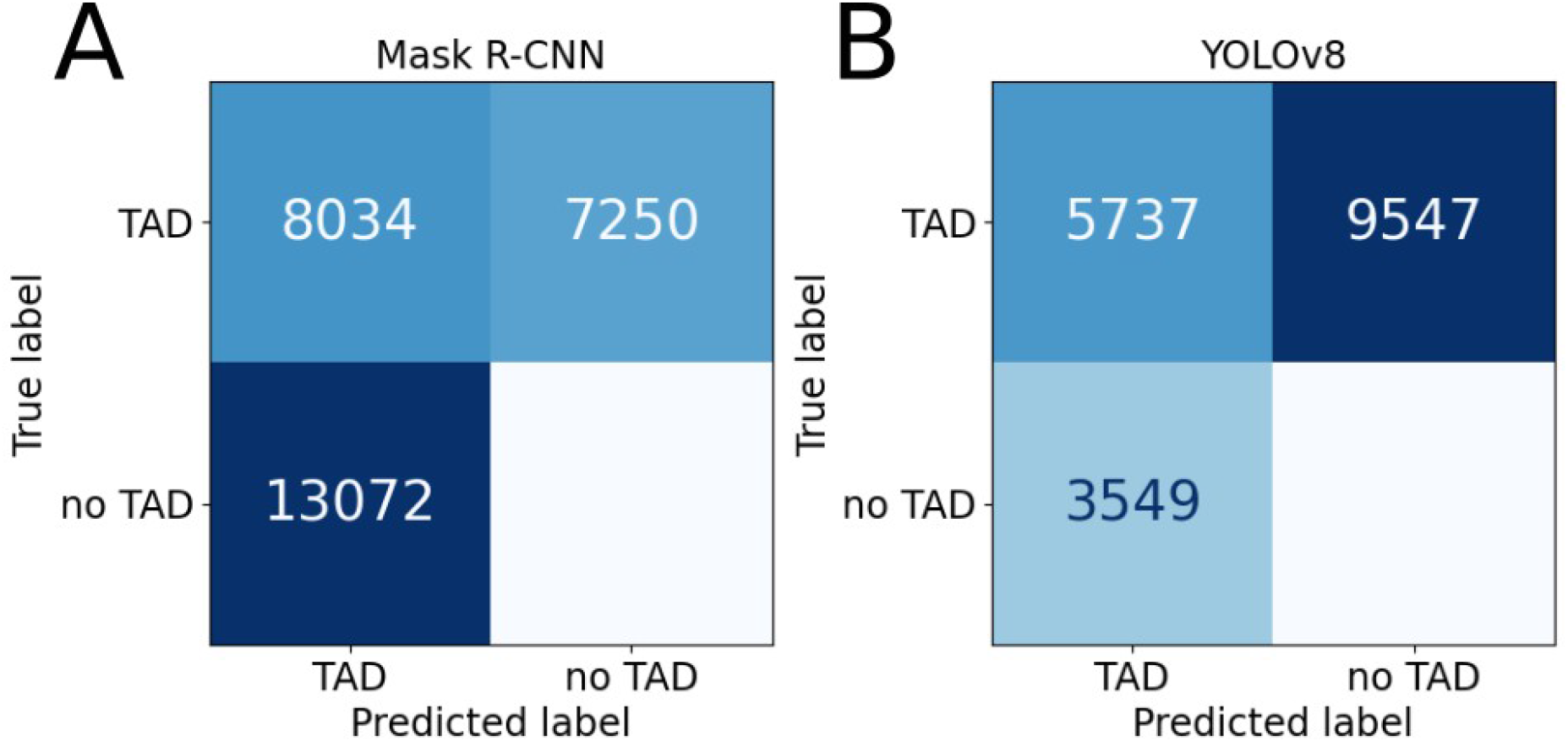
Overlap of TADs detected by YOLOv8 and Mask R-CNN with reference TADs. Confusion matrices for Mask R-CNN (A) and YOLOv8 (B) detected in the HepG2 dataset. A TAD is defined as TP if the maximum IoU with a reference TAD is ≥ 0.8.

**Figure S9:**
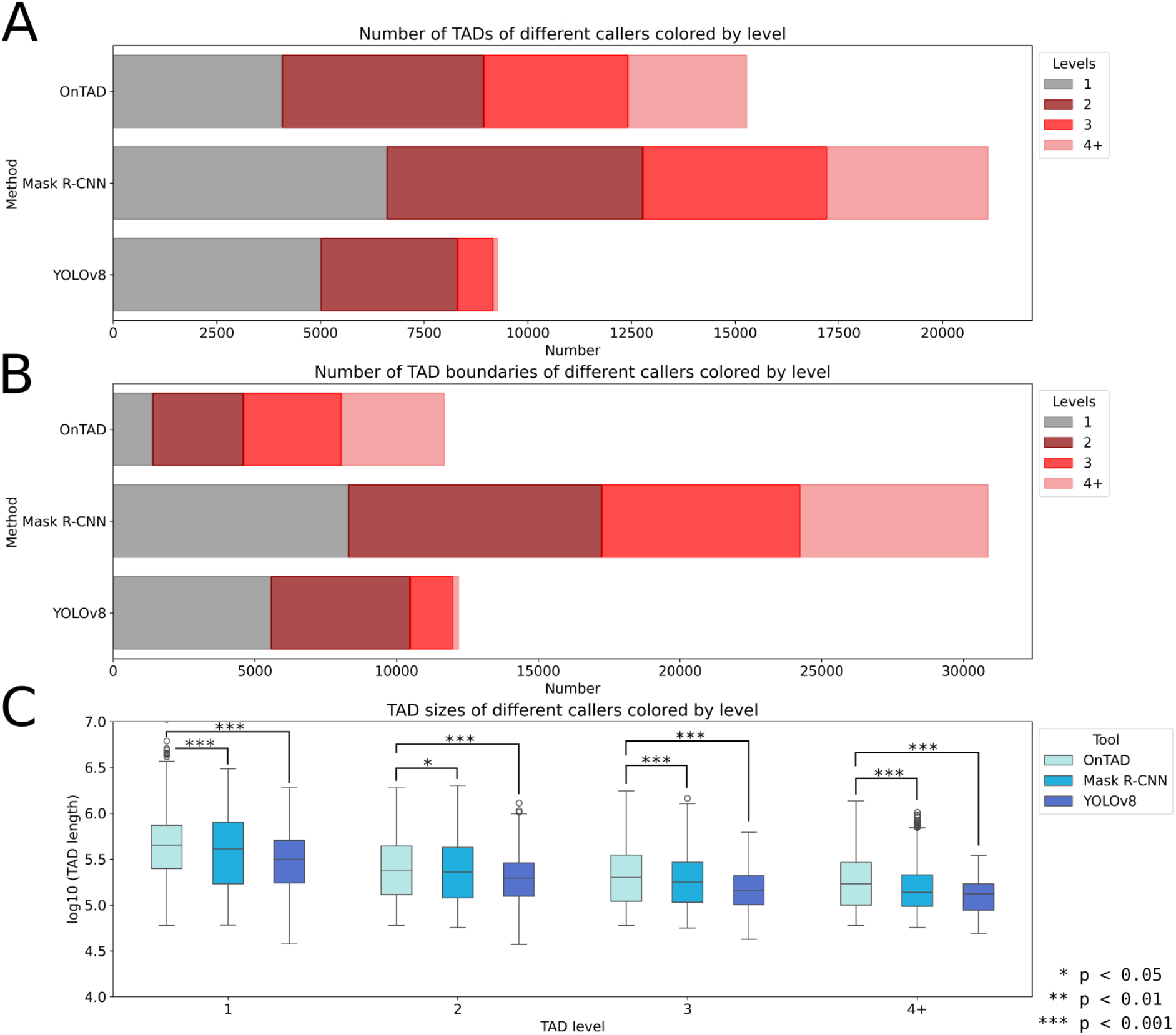
Characteristics of reference TADs, TADs detected by Mask R-CNN and YOLOv8 in the training data. TADs are subdivided according to their nesting level (TADs with a nesting level ≥ 4 are summed in the “4+” group). A shows the number of detected TADs. B shows the number of unique boundaries of TADs subdivided based on TAD level: If one genomic position is the boundary of multiple TADs, we select the highest level of the TADs for B. C shows the sizes of TADs. Significance levels of differences in TAD sizes between reference and both models were calculated using the two-sided Mann-Whitney U test and p-values were adjusted for multiple testing by Benjamini-Hochberg.

**Figure S10:**
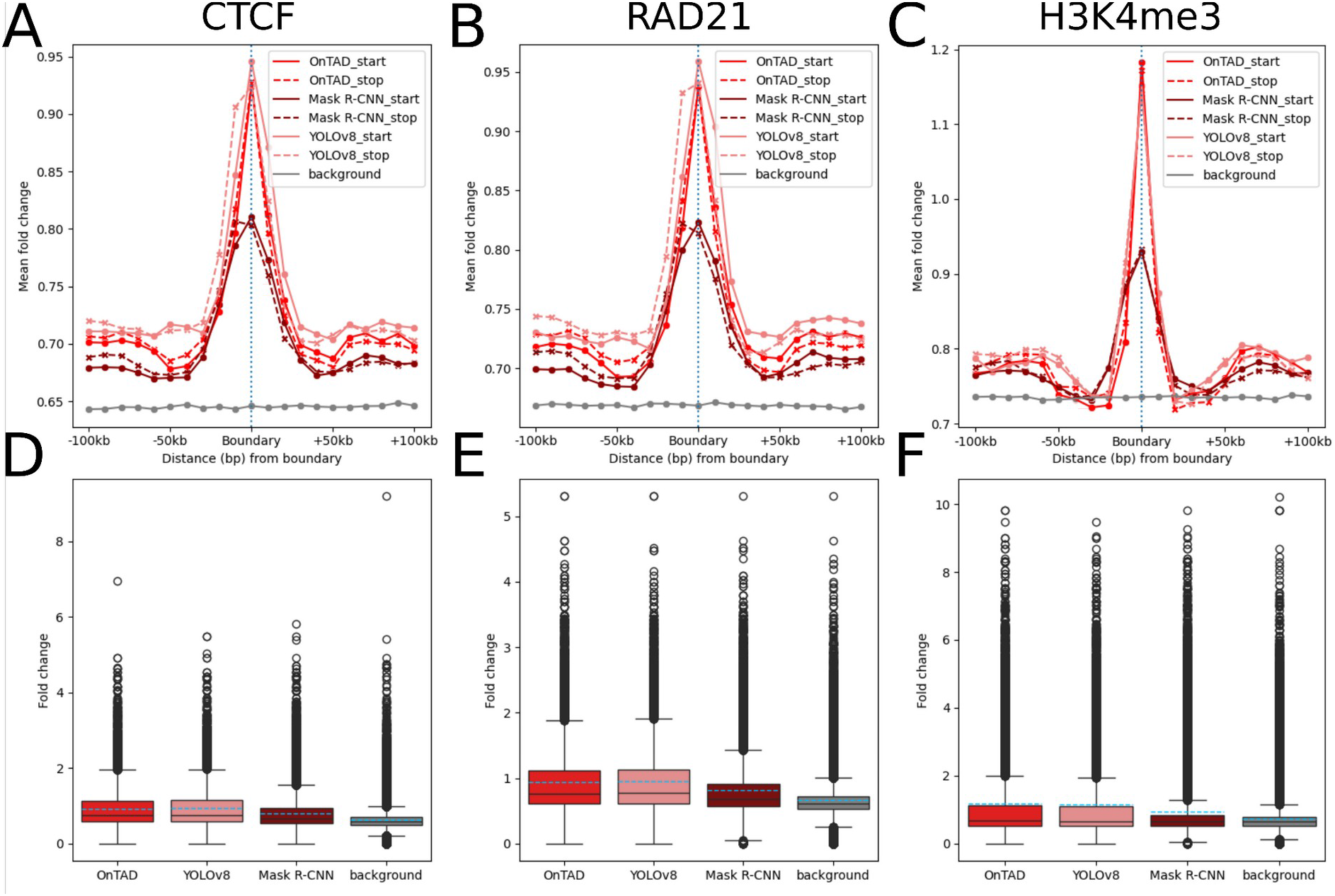
ChIP-Seq signal of CTCF (A, D), RAD21 (B, E) and H3K4me3 (C, F) at TAD boundaries (excluding duplicates) of reference TADs, TADs predicted by YOLOv8 and Mask R-CNN on the HepG2 data used for training. In A, B and C, the averaged fold-change in a 200 kbp window around the boundary (21 bins) is shown to display differences in ChIP-Seq signal. The background signal of randomly drawn positions is added in grey. Blacklist regions (Amemiya et al., 2019) are excluded from background calculation. The position of the predicted boundary is shown as a blue dotted line. In D, E and F, boxplots show the ChIP-Seq signal of boundaries (excluding duplicates). Dashed lines mark the arithmetic means, solid lines the medians. Grey boxes show the background signal of randomly drawn positions.

**Figure S11:**
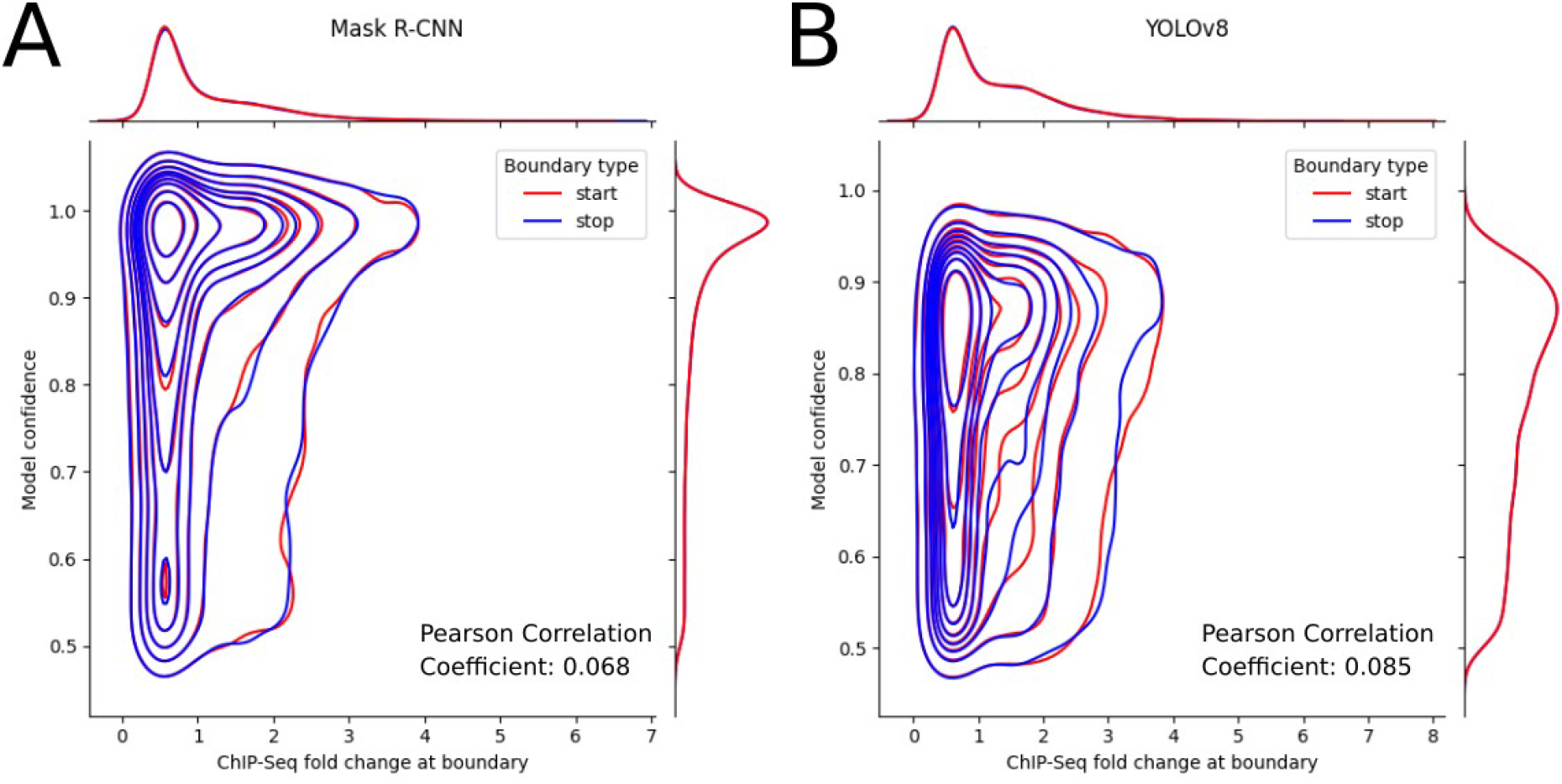
Kernel density estimation of model confidence score and CTCF ChIP-Seq fold change on the boundaries of TADs detected by Mask R-CNN (A) and YOLOv8 (B) in the test dataset. The ChIP-Seq signal is split into the start boundary (red) and stop boundary (blue). Separate kernel density estimations of confidence and CTCF ChIP-Seq fold-change are shown on the right and upper site of the plots. Pearson correlation coefficients of model confidence and ChIP-Seq signal are shown in the lower right.

### Supplementary Methods

#### Comparison of TAD Callers

One of the earlier methods, designed for the identification of unnested and non-overlapping TADs, is TopDom (Shin et al., 2016), a deterministic and window based approach that first computes the average contact frequency among pairs of chromatin regions, then detects TAD boundaries as local minima of this signal, and finally identifies significant domains through statistical testing.

In 2019, OnTAD (An et al., 2019) was published as an efficient method for the identification of hierarchically nested but strictly non-overlapping TADs. In the first step, candidate TAD boundaries are predicted using an adaptive local minimum search algorithm based on TopDom. In the second step, boundaries are partitioned into TADs by recursively splitting the input matrix X_a,b_ in two submatrices X_a,i_ and X_i,b_ where X_i,b_ is the best rightmost TAD based on OnTADs own scoring function.

Recently, Shen et al. introduced HTAD (Shen et al., 2024), a discriminative active learning based approach that enables the iterative incorporation of human feedback (human-in-the-loop strategy). HTAD identifies initial TADs by using a simplified Directionality Index, extracts features from them on Hi-C matrices and uses a binary classifier neural network to identify correct TADs. Resulting TADs can be gapped, nested and overlapping.

Another machine learning based method is TOAST (Gong et al., 2023) utilizing graph auto-encoders and clustering techniques for non-hierarchical TAD recognition where the contact matrix is considered as a graph’s adjacency matrix and the auto-encoder provides embeddings considered as features in later analysis. HDBSCAN (Campello et al., 2013) is used for clustering genomic bins based on their features and bins with the same label are defined to form a TAD. For an extensive list of TAD callers and the types of TAD interactions they detect see (Dali & Blanchette, 2017; Sefer, 2022; Zufferey et al., 2018)

#### YOLO

The first model we evaluate is YOLO, a convolutional neural network architecture, which detects objects in a single step. The image is fed into the fully convolutional part of the model, processed and split into a desired grid size. For each grid cell the model predicts a predefined number of box coordinates (upper left x-coordinate, upper left y-coordinate, height, width), a single confidence score, which corresponds to the overlap with a ground truth box and a number of conditional confidences for each class under the assumption that the bounding box overlaps with an object (Redmon et al., 2015).

YOLO is constantly under development and new YOLO versions are released approximately every year. We used the pretrained YOLOv8 release provided by ultralytics, who maintain and distribute YOLO releases (Jocher et al., 2023).

The YOLOv8 losses include:

- Box-loss (Loss_box_reg) - penalizes incorrect positioning of bounding boxes
- Classification-loss (Loss_classifier) - penalizes incorrect confidence scores
- Distribution focal loss (Loss_dfl) - penalizes incorrect detection of hard-to-detect instances (X. Li et al., 2020)

The names in brackets correspond to the legends of the loss plots in Figure S2.

#### Mask R-CNN

Mask R-CNN – in contrast to YOLO – is a two stage object detection model. The two steps are carried out by two separate models. The first model takes three channel images as input and uses a series of shrinking operations, to continuously reduce the image size. In a shrunken image, each pixel is treated as the center of a set of anchor boxes, which differ by their sizes and aspect ratios. For each anchor box, the model predicts refined bounding box coordinates and a confidence score, which is used to determine if a real object overlaps with the bounding box. The model returns a set of bounding boxes which contain potential object instances and is therefore called region proposal network (RPN) (Ren et al., 2015). Since the objects we want to detect are squares, we use a custom set of anchor boxes with an aspect ratio of 1.

The second part of Mask R-CNN takes the regions of interest in the form of stacked images as input, and predicts a final refinement of the bounding box coordinates, a binary segmentation mask for each object instance and confidence scores for all available classes (He et al., 2017).

To turn the region proposals of the RPN into many small images for the second model, the bounding boxes detected by the RPN need to be extracted from the input image and resized to a predefined size. For this step the model uses ROIalign, a method which resizes a bounding box by placing a grid of the desired size on the box, interpolating new points centered on each grid cell and pooling these new points (He et al., 2017). We use an untrained Mask R-CNN architecture provided by PyTorch (Paszke et al., 2019), which consists of a basic Mask R-CNN architecture in combination with a ResNet-50-FPN backbone from Y. Li et al. (2021).

Since Mask R-CNN is a two stage predictor, both models come with individual sets of loss functions. The RPN losses include:

- Objective-loss (Loss_rpn_objectness) - penalizes wrong confidence scores (object vs background)
- Box-loss (Loss_rpn_box_reg) - penalizes incorrect positioning of bounding boxes

The losses of the second model include:

- Classification-loss (Loss_classifier) - penalizes wrong class-confidence-scores
- Box-loss (Loss_box_reg) - penalizes incorrect refinement of RPN bounding boxes
- Mask-loss (Loss_mask) - penalizes incorrect segmentation masks

The names in brackets correspond to the legends of the loss plots in Figure S2.

#### Selection of data augmentations and thresholds

During training, we apply different data augmentations on the Hi-C matrices and the contained TADs to add more variety to the data. For YOLOv8, we use the default image augmentations provided by the ultralytics training pipeline (Jocher et al., 2023). These augmentations include horizontal and vertical image movement, image scaling, image flipping, merging of four images, erasing of image regions and modification of hue, saturation and brightness. It is important to note that none of these augmentations violates the shape of TADs like e.g. random degree rotation would. Since Mask R-CNN does not have a predefined train-pipeline, we selected the set of image augmentations. They include random rotation by 180 degrees, random cropping centered on the main diagonal, injection of gaussian noise and blur and modifications to brightness, hue and sharpness.

We apply a classification threshold of 0.5 to model detections since it is the midpoint of our classification value range, which reaches from 0 to 1. For NMS filtering, we decided to use an established pairwise IoU threshold of 0.5 (Yang et al., 2026).

### Supplementary Results

#### Further evaluation of TADs detected in the H1-hESC dataset

When comparing reference TADs with TADs detected by both models after confidence- and NMS-filtering in the test dataset, we see that Mask R-CNN detects 26.4% more TADs compared to the reference (Tab. S1A, Fig. S7A). YOLOv8 in contrast detects 25.8% less TADs. Additionally, all three methods detect more TADs in the test dataset than in the training data and the relative increase of TAD numbers increases with higher nesting level: When we compare the number of detected level 1 TADs in both datasets, the number of reference TADs increases by 0.8%, Mask R-CNN detects 3.7% more TADs and YOLOv8 detects 31.7% more TADs (Tab. S1A, Tab S2A). When we compare detected level 4+ TADs between datasets, the observed trend intensifies as the reference TADs increase by 71.6%, Mask R-CNN detects 50.7% more TADs and YOLOv8 detects 228% more TADs.

Since a single bin in Hi-C contact matrices can be the start or stop of multiple nested or adjacent TADs, we also compared the numbers of unique boundaries between the reference, YOLOv8 and Mask R-CNN in the test dataset (Fig S7B, Tab. S1B). If one boundary is shared by multiple TADs, we assign the highest nesting level of the TADs to this boundary. When comparing Figure S7A and B, we see that the reference TADs have the fewest unique boundaries per TAD, followed by YOLOv8 and Mask R-CNN.

Further, we compared the sizes of TADs depending on their nesting level between the three methods (Fig. S7C, Tab. S1C). Reference TADs are the largest at all four nesting levels followed by Mask R-CNN and YOLOv8. We found that TAD sizes of Mask R-CNN and YOLOv8 are always significantly different to reference TADs of the same nesting level.

#### Separation of detected TADs into the fractions *present*, *intermediate* and *new*

In general, TADs detected by YOLOv8 and Mask R-CNN have a high IoU with reference TADs (Fig. S3A). However, since both models detect many FPs, we used the IoU to split TADs detected by the two models into partially overlapping with reference and not overlapping: We label a TAD as *present* if the maximum IoU with a reference TAD is ≥ 0.8 (same definition as TP above), as *new* if the maximum IoU is < 0.2 and as *intermediate* if the IoU is between 0.8 and 0.2. The *new* group contains TADs which were not detected by the reference. The *intermediate* group contains TADs, which cannot be added to one of the two other groups.

The results show that the two models differ in the proportion of *new* and *present* TADs: In the test dataset 8885 TADs (59.1%) of YOLOv8 detections are *present*, while 1597 TADs (10.6%) are *new* (Fig. S3B, Tab. S1E). Of all Mask R-CNN TADs, 11282 TADs (44.0%) are *present* and 4070 TADs (15.9%) are *new*.

#### Evaluation of TADs detected in the HepG2 dataset

Based on the TP-, FP- and FN-TADs detected in the training dataset shown in Figure S8, Mask R-CNN has a TPR of 0.526 and an FNR of 0.474. YOLOv8 in contrast has a TPR of 0.375 and an FNR of 0.625. Mask R-CNN detects 13072 FPs, which is more than 3-fold the number of FPs detected by YOLOv8.

Compared to reference TADs we see that YOLOv8 detects 39.2% less TADs and Mask R-CNN detects 38.1% more TADs. Most of the reference TADs as well as TADs detected by both models have a low nesting level of 1 and 2 followed by 3 and 4+ (Fig. S9A, Tab S2A).

The number of unique TAD boundaries detected by YOLOv8 decreases with increasing nesting level, the number of reference TADs shows the opposite trend and increases with increasing nesting level (Fig. S9B, Tab S2B). In Mask R-CNN detections most of the boundaries are level 2, followed by level 1, level 3 and level 4+.

We also compared the sizes of TADs depending on their nesting level between the three methods in the training data (Fig. S9C, Tab. S2C). Reference TADs are the largest in all four nesting levels followed by Mask R-CNN and YOLOv8. We found that TAD sizes of Mask R-CNN and YOLOv8 are always significantly different to reference TADs of the same nesting level (Fig. S9C).

In the training data, all methods show ChIP-Seq fold-change peaks on TAD boundaries for CTCF, RAD21 and H3K4me3 (Fig. S10), but they are less steep and the fold change on the boundary is lower compared to the test dataset.

In the training data, 61.8% of YOLOv8 TADs are *present* and 7.6% are *new*. Of all Mask R-CNN TADs, 38.1% are *present* and 21.1% are *new*. YOLOv8 detects 37.5% of reference TADs in the training data, while Mask R-CNN detects 52.6%. The corresponding TAD numbers are shown in Tab. S2E.

In the training data 17122 of 21106 TADs (81.1%) detected by Mask R-CNN overlap resulting in 20322 overlaps while only 2494 of 9286 TADs (26.9%) detected by YOLOv8 overlap, resulting in 1631 overlaps.

#### Correlation of model confidence and ChIP-Seq signal in the H1-hESC data

Since YOLOv8 and Mask R-CNN return confidence scores for every detected TAD, we tested if TADs in the test dataset with a higher confidence score show higher CTCF ChIP-Seq signal on their boundaries. Figure S11 shows a fraction of TADs with high confidence score and high ChIP-Seq signal but the overall pearson correlation between model confidence and ChIP-Seq signal is only 0.085 (YOLOv8) and 0.068 (Mask R-CNN).

#### ChIP-Seq signal on boundaries of overlapping TADs in the H1-hESC data

To ensure, that the fraction of overlapping TADs does also show ChIP-Seq signal peaks at the boundaries (which we expect after looking at the signal of all TADs as well as the *new* and *present* fractions) we compared the ChIP-Seq signal on the boundaries of all overlapping TADs as well as TADs with overlaps ≥ 5 bins, which are more likely to be real overlaps not caused by positioning errors (Fig. S6).

The entirety of overlapping TADs as well as the TADs with bigger overlaps (≥ 5 bins) show peaks for all ChIP-Seq signals, however there is only a minor difference between both groups and the fraction containing all overlaps consistently shows slightly higher ChIP-Seq peaks (Fig. S6). Compared to Figure 4, the fold change on the boundaries of overlapping TADs is located between the groups *present* and *new* in Mask R-CNN detections and more towards *new* in YOLOv8 detections. Interestingly, overlapping YOLOv8 TADs show an increase in H3K4me3 signal towards the outside of the TAD.

## Notes

https://doi.org/10.5281/zenodo.23389677

### Competing Interest Statement

The authors have declared no competing interest.

